# Behavioral decomposition reveals rich encoding structure employed across neocortex

**DOI:** 10.1101/2022.02.08.479515

**Authors:** Bartul Mimica, Tuçe Tombaz, Claudia Battistin, Jingyi Guo Fuglstad, Benjamin A. Dunn, Jonathan R. Whitlock

**Affiliations:** Princeton Neuroscience Institute, Princeton University, Princeton, NJ 08544, USA; Kavli Institute for Systems Neuroscience, Norwegian University of Science and Technology, NO-7489 Trondheim, Norway; Department of Mathematical Sciences, Norwegian University of Science and Technology, NO-7489 Trondheim, Norway

**Keywords:** 3D tracking, behavior, posture, rat, electrophysiology, neocortex

## Abstract

The Cortical population code is pervaded by activity patterns evoked by movement, but how such signals relate to the natural actions of unrestrained animals remains largely unknown, particularly in sensory areas. To address this we compared high-density neural recordings across four cortical regions (visual, auditory, somatosensory, motor) in relation to sensory modulation, posture, movement, and ethograms of freely foraging rats. Momentary actions, such as rearing or turning, were represented ubiquitously and could be decoded from all sampled structures. However, more elementary and continuous features, such as pose and movement, followed region-specific organization, with neurons in visual and auditory cortices preferentially encoding mutually distinct head-orienting features in world-referenced coordinates, and somatosensory and motor cortices principally encoding the trunk and head in egocentric coordinates. The tuning properties of synaptically coupled cells also exhibited connection patterns suggestive of different uses of behavioral information within and between regions. Together, our results speak for detailed, multi-level encoding of animal behavior subserving its dynamic employment in local cortical computations.

## Introduction

Much has been learned from studying sensory and motor cortical systems in isolation (Niell 2015; Ebbesen and Brecht 2017; Glickfeld and Olsen 2017; Peters et al. 2017; Nelken 2020), using laboratory tasks in which animals perform *a priori* defined subsets of behaviors in response to experimenter-defined stimuli (Gomez-Marin et al. 2014). However, while such approaches bring essential reliability and control, they restrict the scope of actions animals can express, which limits understanding of the wider array of features to which neural systems respond when animals engage in natural behaviors (Anderson and Perona 2014; Datta et al. 2019; McCullough and Goodhill 2021). This knowledge gap is underscored by observations in head-fixed animals showing that self-generated movements, independent of behavioral tasks, profoundly influence cortical activity patterns (Musall et al. 2019; Salkoff et al. 2019), including in primary visual cortex, with animals in darkness and under no explicit cognitive burden (Stringer et al. 2019). Recent work has started illuminating the role of behavioral modulation in different sensory cortices (Meyer et al. 2018; Schneider et al. 2018; Vélez-Fort et al. 2018; Stringer et al. 2019; Bouvier et al. 2020; Guitchounts et al. 2020b; Parker et al. 2020; Schneider 2020), but such systems are rarely studied in tandem, and often restricted to actions expressed under heavy experimental constraints. Consequently, it is not well understood how movement-driven signals in cortex reflect the animals’ natural behavioral repertoire, nor is it known if the features encoded vary from region to region, for example, to support different types of sensory processing.

It is becoming possible to address such questions in freely behaving animals, owing to advances in quantitative pose estimation (Mathis et al. 2018; Pereira et al. 2018; Dunn et al. 2021; Marshall et al. 2021b), as well as unsupervised machine learning approaches (Berman et al. 2014; Wiltschko et al. 2015; Marshall et al. 2021a) for classifying unique actions based on underlying structure in tracking data. When paired with neural recordings, such techniques have afforded a range of recent discoveries, including how subcortical circuits generate sub-second patterns of behavior (Markowitz et al. 2018) or encode action space (Klaus et al. 2017), the characterization of escape behaviors (Storchi et al. 2020), or how different pharmacochemical substances leave tractable traces on the behavioral landscape (Wiltschko et al. 2020). Furthermore, machine learning has also enabled researchers to infer rodents’ emotional states from facial videos (Dolensek et al. 2020) or control virtual rodent behavior (Merel et al. 2019), demonstrating the full promise in carefully quantifying animal actions.

Here, to better understand how behavioral representations might be differentially employed across cortical systems, we combined detailed quantification of the behavior of unrestrained rats at multiple levels, from naturalistic actions to elementary poses, with dense single unit electrophysiology in four separate sensory and motor cortical areas. We found that all cortical networks encoded both higher- and lower-level behavioral representations, but only the latter were characterized by distinct, subregion-specific population codes, possibly utilized for locally-relevant computations, underscoring the importance of multilevel analysis of naturalistic actions.

## Results

We combined 3D motion capture with chronic Neuropixels recordings to track the heads and backs of freely foraging rats while recording large ensembles of single units from primary motor and somatosensory cortices (**Fig. 1A**, top; 4 animals, 1,532 and 792 cells, respectively; **methods**) or, visual and auditory cortices (**Fig. 1A**, bottom right; 3 animals, 1,633 and 526 cells, respectively). Recording sites were localized to different cortical regions using a custom pipeline (**fig. S1A-E** and **methods**) which allowed us to triangulate the anatomical position of individual channels along each probe (**fig. S1F-G**). Single units were classified by spiking profile (**fig. S2** and **methods**) as well as specific sub-regions where they were recorded (*e.g.*, S1 hindlimb region, S1 trunk, primary or secondary auditory, *etc.*) (Paxinos and Watson 2007) in four overarching areas (motor, somatosensory, visual, auditory).

**Fig. 1.**
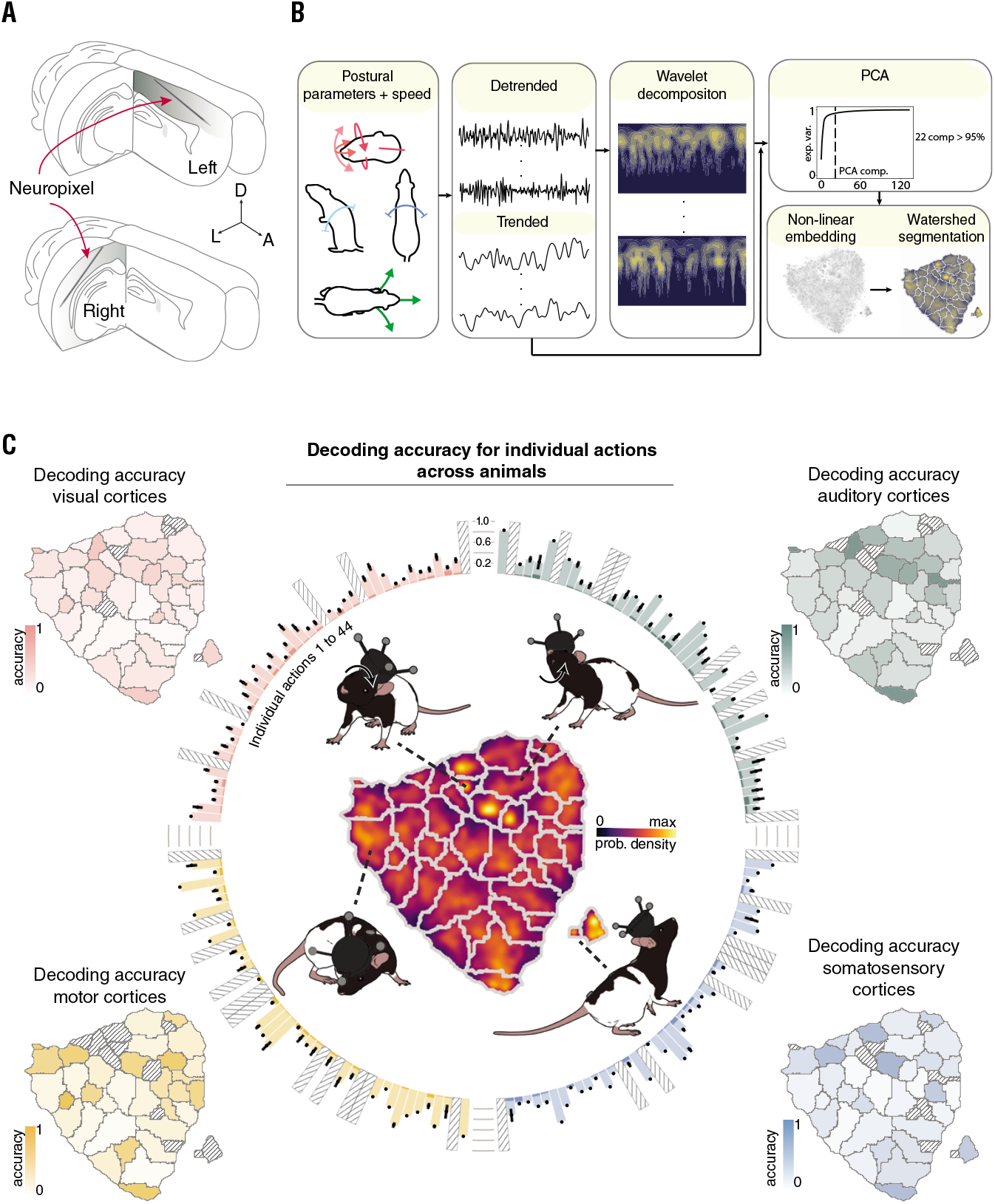
Ensemble decoding of natural actions is robust in visual, auditory, motor and somatosensory cortices. **(A)** (Top) Neuropixels probes implanted in the left hemisphere were tilted 60-70° in the sagittal plane to travel anteriorly from primary somatosensory to primary motor cortex. (Bottom) Separate animals were implanted with probes in the right hemisphere tilted 45-50°in the coronal plane to progress laterally through primary and secondary visual and auditory cortices. **(B)** (Left) To extract discernible actions we first generated time series for each postural feature of the head (pitch, azimuth, roll) and back (pitch, azimuth), together with running speed and neck elevation. (Middle) The data were then detrended and decomposed spectrally using a Morlet wavelet transform. (Right) Features from all animals were sub-sampled and co-embedded in two dimensions using t-SNE, followed by watershed segmentation, producing a map in which distinct, identifiable actions were separated into discrete categories. **(C)** (Middle) A map of distinct actions, color-coded for occupancy, shows the cumulative sampling for each of the 44 actions pooled across all animals; the circumferential bar graph shows the mean decoding accuracy for each action in visual (pink), auditory (cyan), motor (yellow) and somatosensory (blue) cortices (error bars denote ±SEM across different rats and darker colors signify the performance of the decoder consisting only of a prior distribution; actions with insufficient sampling are indicated by striped bars). Four separate examples of actions decoded from each region are shown around the map. Mean decoding accuracy across animals is depicted for each action in the outer t-SNE maps, color-coded by cortical area; actions with insufficient sampling are filled with stripes.

In a given environment, animal behavior can be assessed at different levels of complexity, ranging from elementary poses to identifiable, species-typical actions like rearing, running or turning (Gris et al. 2017). To isolate instances at the higher end of that hierarchy, we drew upon existing approaches (Berman et al. 2014; Marshall et al. 2021a) to transform raw tracking data pooled across rats into sets of postural and movement features, and ultimately into a time-resolved ethogram (**Fig. 1B** and **methods**). The animals’ combined ethogram consisted of 44 independent modular actions (**Fig. 1C, fig. S3A** and **B**) identifiable by dissimilarly evoked rudimentary features (**fig. S4A** and **B**), whose sequence order was best described by a set of transition probabilities (**fig. S3C**). For example, the action “running, head up” was characterized by higher speeds than “running, head level, scanning”, and was most often followed by the latter. As expected with freely behaving rodents (Wiltschko et al. 2015; Marshall et al. 2021a), actions comprising the ethogram were not sampled equally (**Fig. 1C**, center) and, although the animals’ behavior varied across sessions (**fig. S3D**, top), actions were expressed with greater commonality within individuals than between them (**fig. S3E**, top).

With the behavioral phenotype of the rats profiled, we sought to characterize to what degree neural responses in different cortical structures map onto the spectrum of identified actions. We limited the analyses to recordings conducted in darkness in order to minimize visual confounds, and found stable encoding of every considered action by individual neurons in each cortical region (visual, 51%; auditory, 55%; motor, 58%; somatosensory, 56%), with most cells responding to multiple actions, and fewer to single actions (**fig. S3F; movies S1-S6** and **methods**). This was true for all animals, irrespective of the specific anatomical placement of the recording probe, and the distribution of encoded actions was similar across regions (**fig. S3E**, bottom). Crucially, decoding analyses at the ensemble level revealed that nearly all of the 44 independent actions with sufficient sampling could be predicted beyond chance in any of the four overarching areas (**Fig. 1C** and **table 1**). The decoding accuracies from sessions with more than 100 neurons were positively correlated between regions (Spearman’s *ρ*=0.39 0.11 (mean std), *p*=.007; permutation test), underscoring cross-regional similarities in action modulation.

**Table 1.** Mean decoding accuracy for each behavior in each cortical region. The mean and standard deviation of decoding accuracy for each of the 44 actions with sufficient sampling. For each region, decoding accuracy rates for actual data are in the left column and shuffled data are in the right column.

| Decoding accuracy (mean ± std) |  |  |  |  |  |  |  |  |  |
| --- | --- | --- | --- | --- | --- | --- | --- | --- | --- |
| Insufficient sampling<br>//////////////// |  |  |  |  |  |  |  |  |  |
| # | Actions | visual | shuffled | auditory | shuffled | motor | shuffled | somatosensory | shuffled |
| 1 | Still, sitting | 0.58 ± 0 | 0.01 ± 0 | 0.84 ± 0 | 0.01 ± 0 | 0.74 ± 0 | 0.01 ± 0 | //////////////// | //////////////// |
| 2 | Rearing, pre-rearing | 0.47 ± 0.1 | 0.03 ± 0.01 | //////////////// | //////////////// | //////////////// | //////////////// | 0.54 ± 0 | 0.02 ± 0 |
| 3 | Turn left | 0.18 ± 0.04 | 0.02 ± 0.01 | 0.22 ± 0.03 | 0.02 ± 0.01 | 0.3 ± 0.13 | 0.02 ± 0.01 | 0.28 ± 0.01 | 0.02 ± 0 |
| 4 | Walk, CW head roll | 0.21 ± 0.05 | 0.02 ± 0.01 | 0.35 ± 0.06 | 0.03 ± 0.01 | 0.61 ± 0.03 | 0.02 ± 0.01 | 0.11 ± 0 | 0.02 ± 0 |
| 5 | Still, head down left, curl left | 0.2 ± 0.04 | 0.06 ± 0.01 | 0.33 ± 0.14 | 0.06 ± 0.01 | 0.25 ± 0.06 | 0.06 ± 0.01 | 0.22 ± 0.05 | 0.05 ± 0.01 |
| 6 | Walk, head left | 0.42 ± 0.06 | 0.01 ± 0 | 0.78 ± 0 | 0.02 ± 0 | 0.41 ± 0 | 0.01 ± 0 | //////////////// | //////////////// |
| 7 | Still, eating, back hunched | //////////////// | //////////////// | //////////////// | //////////////// | 0.53 ± 0 | 0.01 ± 0 | 0.43 ± 0 | 0.01 ± 0 |
| 8 | Still, head slight down left | 0.33 ± 0.06 | 0.03 ± 0.01 | 0.52 ± 0 | 0.02 ± 0 | 0.76 ± 0.05 | 0.03 ± 0.01 | 0.35 ± 0.13 | 0.02 ± 0 |
| 9 | Walk, head bobble | 0.22 ± 0.07 | 0.02 ± 0.01 | 0.48 ± 0.07 | 0.03 ± 0.01 | 0.47 ± 0.01 | 0.02 ± 0 | 0.53 ± 0.02 | 0.02 ± 0 |
| 10 | Walk, head up left | 0.13 ± 0.03 | 0.02 ± 0.01 | 0.19 ± 0.05 | 0.03 ± 0 | 0.17 ± 0 | 0.02 ± 0 | 0.14 ± 0 | 0.02 ± 0 |
| 11 | Walk, head level, slight rightward movement | 0.26 ± 0.07 | 0.02 ± 0 | 0.19 ± 0.07 | 0.02 ± 0 | 0.4 ± 0 | 0.01 ± 0 | //////////////// | //////////////// |
| 12 | Waddle, head left | 0.06 ± 0 | 0.01 ± 0 | //////////////// | //////////////// | //////////////// | //////////////// | //////////////// | //////////////// |
| 13 | Still, head down left | //////////////// | //////////////// | //////////////// | //////////////// | //////////////// | //////////////// | //////////////// | //////////////// |
| 14 | Walk, head scanning | 0.08 ± 0.04 | 0.05 ± 0.01 | 0.18 ± 0.07 | 0.05 ± 0 | 0.22 ± 0.06 | 0.05 ± 0.01 | 0.21 ± 0.04 | 0.06 ± 0.01 |
| 15 | Still, head level, left | 0.27 ± 0.07 | 0.03 ± 0.01 | 0.35 ± 0.1 | 0.02 ± 0 | 0.44 ± 0.11 | 0.02 ± 0.01 | 0.1 ± 0.04 | 0.03 ± 0.01 |
| 16 | Still, head scanning | 0.43 ± 0.08 | 0.01 ± 0 | 0.41 ± 0 | 0.01 ± 0 | //////////////// | //////////////// | //////////////// | //////////////// |
| 17 | Walk, slight head up, CW roll | 0.26 ± 0.04 | 0.04 ± 0.01 | 0.28 ± 0.1 | 0.04 ± 0.01 | 0.2 ± 0.1 | 0.04 ± 0.01 | 0.31 ± 0.04 | 0.04 ± 0 |
| 18 | Still, head slight left | 0.45 ± 0 | 0.01 ± 0 | 0.67 ± 0 | 0.02 ± 0 | //////////////// | //////////////// | 0.17 ± 0 | 0.01 ± 0 |
| 19 | Still, back hunched | 0.2 ± 0.04 | 0.05 ± 0.02 | 0.13 ± 0.05 | 0.05 ± 0.01 | 0.23 ± 0.1 | 0.05 ± 0 | 0.23 ± 0.03 | 0.06 ± 0.01 |
| 20 | Still, CW head roll | 0.33 ± 0.07 | 0.04 ± 0 | 0.51 ± 0.03 | 0.04 ± 0.01 | 0.21 ± 0.04 | 0.04 ± 0 | 0.16 ± 0.02 | 0.05 ± 0.01 |
| 21 | Walk, slight CW head roll, left head azimuth | 0.08 ± 0.02 | 0.05 ± 0.01 | 0.11 ± 0.03 | 0.04 ± 0.01 | 0.2 ± 0.09 | 0.05 ± 0.01 | 0.12 ± 0.01 | 0.03 ± 0 |
| 22 | Running, head level, scanning | 0.25 ± 0.05 | 0.08 ± 0.02 | 0.28 ± 0.1 | 0.08 ± 0.02 | 0.07 ± 0.04 | 0.08 ± 0.01 | 0.3 ± 0.04 | 0.08 ± 0.01 |
| 23 | Running, head up | 0.53 ± 0.08 | 0.03 ± 0.01 | 0.84 ± 0 | 0.03 ± 0 | 0.75 ± 0.12 | 0.03 ± 0 | 0.66 ± 0.06 | 0.03 ± 0 |
| 24 | Still, CW head roll, slight up | 0.35 ± 0.09 | 0.03 ± 0.01 | 0.7 ± 0 | 0.02 ± 0 | 0.66 ± 0.03 | 0.03 ± 0 | 0.72 ± 0 | 0.04 ± 0 |
| 25 | Waddle, head centered | 0.18 ± 0.04 | 0.03 ± 0 | 0.46 ± 0.11 | 0.02 ± 0 | 0.63 ± 0.06 | 0.03 ± 0 | 0.37 ± 0 | 0.02 ± 0 |
| 26 | Walk, CW Headroll | 0.18 ± 0.05 | 0.04 ± 0.01 | 0.39 ± 0.1 | 0.05 ± 0.01 | 0.12 ± 0.04 | 0.05 ± 0.02 | 0.06 ± 0.01 | 0.05 ± 0 |
| 27 | Still, CCW head roll | //////////////// | //////////////// | //////////////// | //////////////// | //////////////// | //////////////// | //////////////// | //////////////// |
| 28 | Still, head level | 0.19 ± 0.01 | 0.03 ± 0 | 0.58 ± 0.06 | 0.02 ± 0.01 | //////////////// | //////////////// | 0.62 ± 0 | 0.02 ± 0 |
| 29 | Walk, head-turn right | 0.1 ± 0.05 | 0.07 ± 0.02 | 0.16 ± 0.07 | 0.08 ± 0.02 | 0.08 ± 0 | 0.07 ± 0.01 | 0.16 ± 0.06 | 0.07 ± 0.01 |
| 30 | Still, head up | 0.22 ± 0.03 | 0.01 ± 0.01 | //////////////// | //////////////// | //////////////// | //////////////// | //////////////// | //////////////// |
| 31 | Head slight up, right | 0.27 ± 0.06 | 0.04 ± 0.01 | 0.43 ± 0.04 | 0.03 ± 0.01 | 0.25 ± 0.09 | 0.04 ± 0 | 0.16 ± 0.02 | 0.03 ± 0 |
| 32 | Walk, head & body turned right | //////////////// | //////////////// | //////////////// | //////////////// | 0.16 ± 0 | 0.01 ± 0 | 0.35 ± 0 | 0.02 ± 0.01 |
| 33 | Still, head right, CW roll, movement | 0.41 ± 0.07 | 0.04 ± 0.01 | 0.41 ± 0.06 | 0.04 ± 0.01 | 0.25 ± 0.07 | 0.04 ± 0.01 | 0.09 ± 0.03 | 0.03 ± 0 |
| 34 | CCW roll, head moving slightly right | 0.57 ± 0.08 | 0.02 ± 0 | 0.86 ± 0 | 0.02 ± 0 | //////////////// | //////////////// | 0.22 ± 0 | 0.02 ± 0 |
| 35 | Still, head right, CW roll | 0.33 ± 0.02 | 0.02 ± 0.01 | 0.18 ± 0 | 0.01 ± 0 | 0.59 ± 0 | 0.02 ± 0 | 0.22 ± 0.02 | 0.02 ± 0 |
| 36 | Walk, slight head down right | 0.09 ± 0.03 | 0.05 ± 0.01 | 0.13 ± 0.05 | 0.05 ± 0.01 | 0.13 ± 0.04 | 0.04 ± 0.01 | 0.08 ± 0.01 | 0.06 ± 0.01 |
| 37 | Still, slight CCW head roll | 0.13 ± 0 | 0.02 ± 0 | //////////////// | //////////////// | //////////////// | //////////////// | 0.38 ± 0 | 0.01 ± 0 |
| 38 | CCW head roll, head down, moving right | 0.29 ± 0.07 | 0.02 ± 0 | 0.43 ± 0.13 | 0.02 ± 0.01 | 0.71 ± 0 | 0.02 ± 0 | 0.65 ± 0 | 0.02 ± 0 |
| 39 | Still, head right | 0.18 ± 0.04 | 0.04 ± 0 | 0.34 ± 0.12 | 0.04 ± 0.01 | 0.18 ± 0.05 | 0.03 ± 0.01 | 0.21 ± 0.04 | 0.03 ± 0.01 |
| 40 | Walk, head right, slight up | 0.19 ± 0.06 | 0.04 ± 0.01 | 0.33 ± 0.06 | 0.04 ± 0.01 | 0.22 ± 0.06 | 0.03 ± 0.01 | 0.26 ± 0.02 | 0.05 ± 0.01 |
| 41 | Still, head turn right | 0.39 ± 0.07 | 0.01 ± 0.01 | 0.29 ± 0.11 | 0.01 ± 0.01 | 0.87 ± 0 | 0.02 ± 0 | 0.14 ± 0 | 0.02 ± 0 |
| 42 | Still, curled right, head down right | 0.17 ± 0.04 | 0.07 ± 0.02 | 0.29 ± 0.11 | 0.07 ± 0.02 | 0.16 ± 0.03 | 0.06 ± 0.01 | 0.14 ± 0.02 | 0.07 ± 0.02 |
| 43 | Still, back hunched, head down | 0.28 ± 0.06 | 0.04 ± 0.01 | 0.23 ± 0.07 | 0.04 ± 0.01 | 0.59 ± 0.1 | 0.04 ± 0 | 0.34 ± 0.12 | 0.04 ± 0.02 |
| 44 | Rearing up | //////////////// | //////////////// | //////////////// | //////////////// | //////////////// | //////////////// | //////////////// | //////////////// |

The widespread representation of momentary actions prompted us to more closely inspect tuning to elementary features of the animals’ ethograms, including posture and movement of the head, neck and trunk (along Euler axes of pitch, azimuth and roll), as well as whole-body movements including self-motion and running speed (23 features in total; **Fig. 2**, bottom and **methods**). Head posture and movement were further distinguished as relative to the back (egocentric) or relative to the world (allocentric). We found reliable neuronal responses to nearly any of the measured features in all areas and animals, similar to those previously reported in motor and posterior parietal cortices (Mimica et al. 2018; Zimmermann et al. 2020) (**fig. S5**). Coding properties of the cells were established by a statistical model selection framework (**fig. S6A** and **methods**), where we again relied principally on recordings conducted in the dark. For each region we quantified (i) the fraction of neurons with selected covariates that provided the single best out-of-sample fit relative to the intercept-only model (**Fig. 2**, inset pie charts), (ii) the proportion to which each covariate was selected among others (**Fig. 2**, polar plot wedge widths), (iii) mean cross-validated relative log-likelihood ratio (rLLR, **methods**) for each selected feature (**Fig. 2**, polar plot wedge heights), (iv) the distribution of covariate counts, *i.e.* model sparsity (**Fig. 2**, gray histograms), and (v) the distribution of mean cross-validated pseudo-R^2^ (methods) values across selected models for each area (**fig. S7**). This breakdown is further graphically clarified on data pooled across regions (**fig. S6B**).

**Fig. 2.**
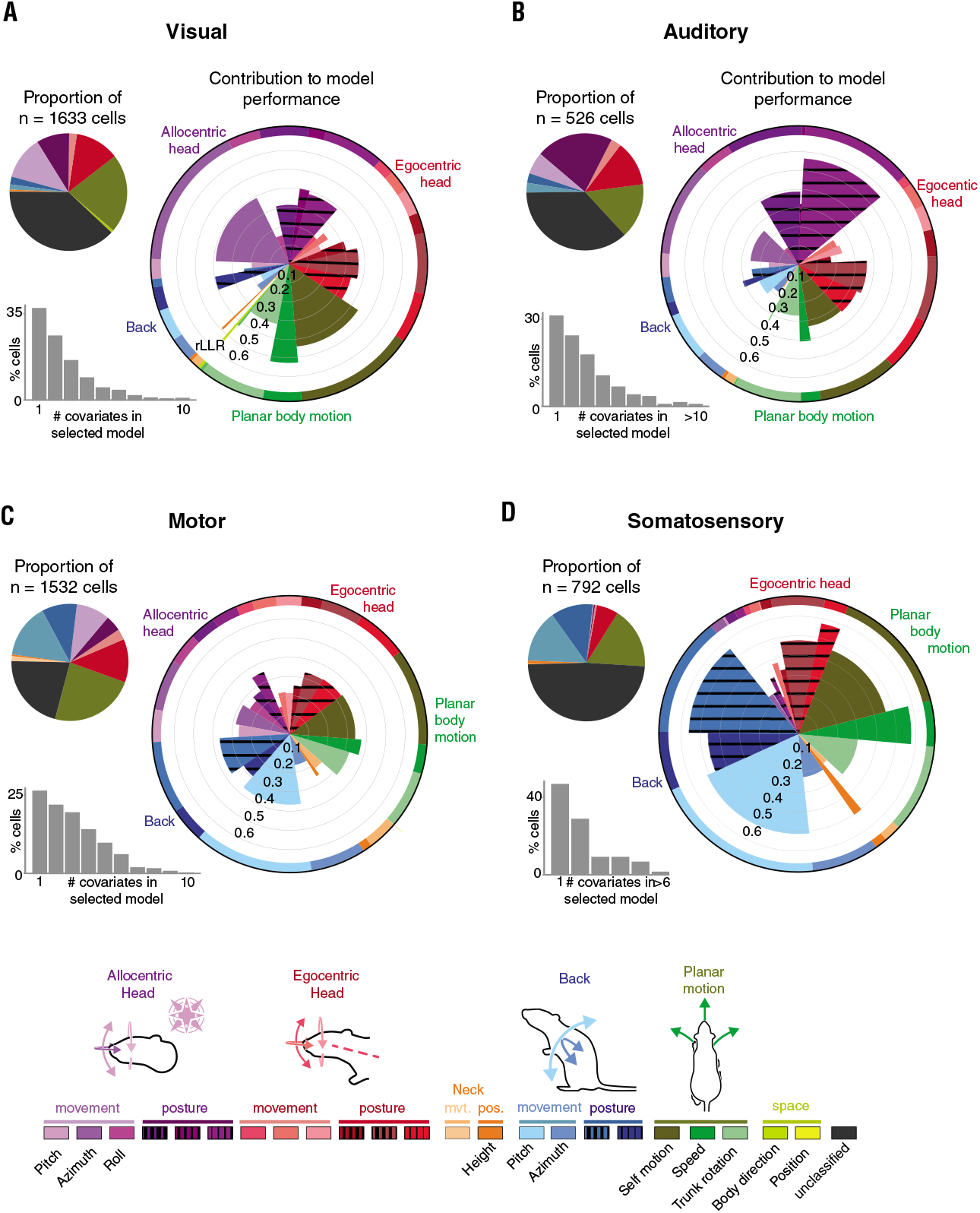
Distinct cortices show differential tuning to posture, movement and self-motion. **(A)** (Top left) The fraction of single units in visual cortices (V1, V2L and V2M) that incorporated specific behavioral features as the first covariate (largest mean cross-validated log-likelihood among single covariates models) in model selection (for feature identification, refer to the color-coded legend at bottom). (Lower left) The percentages of single units statistically linked to one or any larger number of behavioral covariates. (Right) The relative importance of individual covariates in the data, where the width of each wedge reflects the fraction to which each covariate was represented among all those selected, and length denotes the mean cross-validated rLLR (**methods**) of each covariate across the set of models it was included in. **(B)** Same as (A) but for auditory cortices (A1 and A2D). **(C)** Same as (A) but for the primary motor cortex (M1). **(D)** Same as (A) but for the primary somatosensory cortex (S1HL and S1Tr). (Bottom) Two color gradients were used to convey GLM results: one in pie charts (denoted by thick elongated lines) with related features grouped together, and one in polar plots (denoted by clear and striped rectangles) with each representing an individual feature.

The majority of cells in visual (62%) and auditory (63%) cortices encoded at least one behavioral covariate (**Fig. 2A** and **B**, pie and polar charts), which was juxtaposed by a lower rate in somatosensory (51%) and the highest one in primary motor cortex (79%) (**Fig. 2C** and **D**, pie and polar charts). Since anatomical inputs and cortical dynamics vary across layers (Harris and Mrsic-Flogel 2013; Bouvier et al. 2020; Jordan and Keller 2020), we note that single units across visual and auditory cortices were not sampled equally from all layers due to probe implantation angles (V1 (L2/3 to L6), V2L (L6), A2D (L5 and 6) and A1 (L5 and 6; L4 in one animal); **fig. S8A**). Only superficial layers were recorded in somatosensory cortex (S1Tr - L2/3; S1HL - L2/3; not shown), whereas recordings in motor cortex included both superficial and deep layers of (M1 - L2/3 and L5; not shown).

Coding in visual cortex was strongest for combinations of features capturing allocentric head movement along the horizontal plane (specifically, azimuthal head movement and planar body motion; **Fig. 2A** and **table 2**, bottom), and egocentric head posture. The largest fraction of cells in auditory cortex, on the other hand, principally encoded features conveying gravity-relative head orientation (**Fig. 2B**, pie chart), with spiking activity best fit by models for allocentric head roll and pitch, followed by egocentric head posture (**Fig. 2B**, polar chart and **table 2**, bottom). For both auditory and visual cortices, the overall number of tuned cells and the proportion of covariates did not differ considerably between primary and secondary subareas, but neurons in primary areas consistently exhibited higher rLLRs (**fig. S8B**, right), and were better fit by sparser models, which was clearly contrasted by more complex behavioral modulation in secondary cortices (**fig. S8C**). The fact that the Neuropixels shanks spanned several millimeters of tissue allowed us to test for anatomical gradients in features encoded visual and auditory regions (**fig. S9A** and **methods**), which revealed: (1) an increase in allocentric head posture representation as probes progressed from V1 to V2L, peaking in A2D, (*χ*^2^(7)=29.5, p=4.8e-5), (2) a peak in allocentric head movement features in V2L (*χ*^2^(7)=13.09, p=.04), and (3) a strong tendency of planar body motion features (*e.g.*, self-motion and body direction) to group on the border between V2L and deeper layers of V1 (*χ*^2^(7)=18.2, p=.006).

**Table 2.**
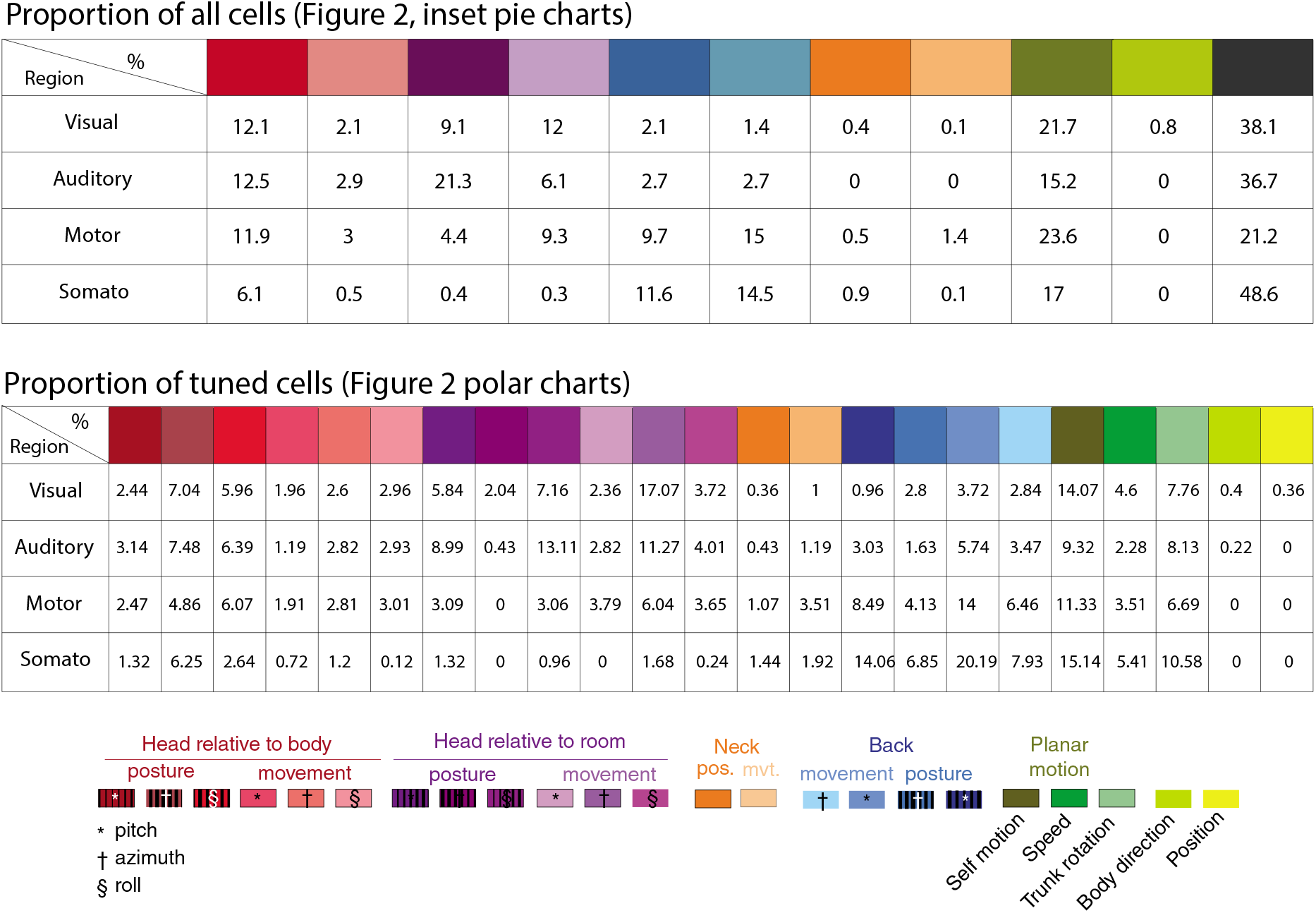
Numerical distribution of behavioral features encoded in each cortical region. (Top) The fraction of single units in visual, auditory, somatosensory and motor cortices that incorporated specific behavioral features as the first covariate (largest increase of the mean cross-validated relative log-likelihood ratio (rLLR) relative to the null-model) in model selection (refer to the color-coded legend at bottom for feature identification). (Bottom) The percentages of single units statistically linked to one or any larger number of behavioral covariates in each cortical region.

Primary motor cortex principally encoded planar body motion, back movement and egocentric head posture (**Fig. 2C**, pie chart and **table 2**), which corresponded well with back movement and self-motion being the most represented features across classified cells (**Fig. 2C**, polar chart). Mean cross-validated rLLRs were moderate and strikingly similar across the most prominent features (**Fig. 2C**, polar plot), in agreement with complex models best accounting for motor cortex spiking activity (**Fig. 2C**, gray histogram). These models also generally exhibited higher explanatory power compared to those selected in other areas (**fig. S7**, median cross-validated pseudo-R^2^ for auditory (.02), visual (.01), motor (.03) and somatosensory (.02) areas). Since we uncovered broad representations of head kinematics, we performed separate recordings with and without a 15 g weight mounted on the animals’ implants (**fig. S10A**, top left) to test whether rate encoding in our data genuinely reflected such kinematics or, rather, muscle forces required to maintain a stable posture (Kakei et al. 1999; Crammond and Kalaska 1996). The added weight caused the animals’ heads to roll slightly towards one side (**fig. S10A**, right), but otherwise had very little effect on behavior or tuning in either motor or visual areas, except for higher firing rates and stability of motor neurons in light sessions for egocentric head azimuth posture (**fig. S10E** and **F, table 3**). Likewise, modeling comparisons between the light and weight sessions of motor cells exhibited no obvious encoding differences (**fig. S10G**).

**Table 3.**
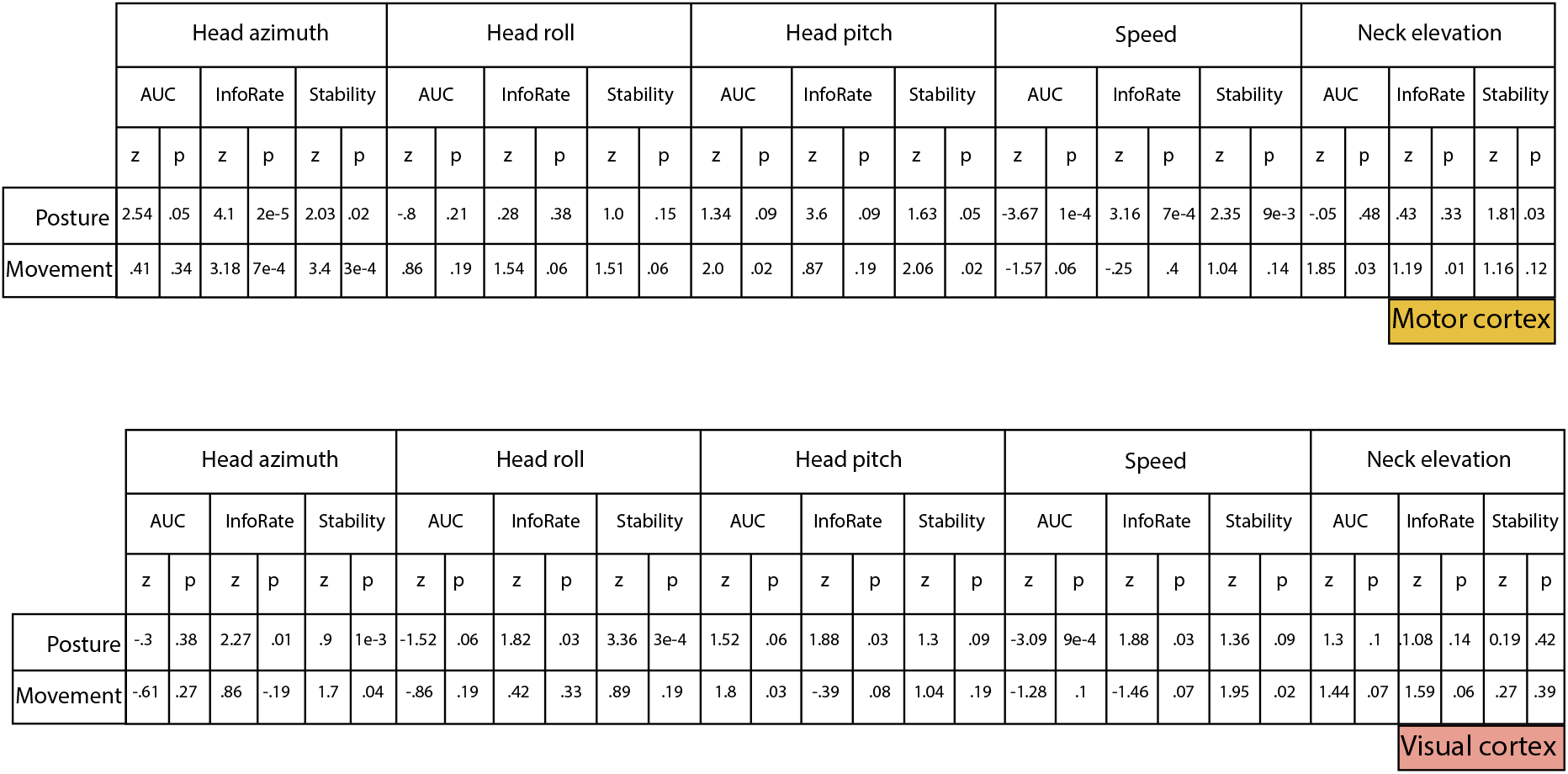
Summary of statistical comparisons between “weight” and “light” sessions. (Top) Details of statistical testing procedures in relation to the area under the curve (AUC), information rate and stability across weight and light sessions in motor cortex as shown in **Fig. S10F**. “z” refers to z-score and “p” refers to p-value in comparison to an empirical shuffle distribution. (Bottom) Same, but for visual neurons.

Finally, somatosensory cortex primarily responded to planar body motion, back movement and posture, but also had the largest proportion of unclassified units (**Fig. 2D**, pie chart). Compared to other areas, somatosensory neurons were better fit by sparser models, containing mainly either one or two covariates (**Fig. 2D**, gray histogram). Such models were largely dominated by back postural and movement features (**Fig. 2D**, polar plot). Topographical organization of features was also assessed across somatomotor cortices, revealing two rostrocaudal gradients: (1) egocentric head posture and movement encoding neurons were more frequently found in rostral M1 than caudal motor or somatosensory areas (posture: *χ*^2^(7)=37.7, p=1.3e-6; movement: *χ*^2^(7)=106.9, p=8.8e-21), (2) back representations dominated in S1HL and caudal motor areas, but much less so in rostral M1 (posture: *χ*^2^(7)=18.8, p=.004; movement: *χ*^2^(7)=41.02, p=2.9e-7) (**fig. S9B**). Having also captured head kinematics with inertial measurement units (IMUs), we compared modeling results based on IMU-generated covariates to those based on optical tracking. Although both approaches revealed encoding of allocentric head features in auditory and visual cortices, only optical tracking, which included tracking of the back, was suited to distinguish the dominant egocentric head and back-related receptivity in motor and somatosensory areas (**Fig. 2A-D; fig. S11**).

To quantify the overlap of behavioral and sensory modulation in visual and auditory regions we conducted recordings with the room lights on and off and, separately, with intermittent presentation of 5s white noise sequences (**Fig. 3A-C** and **fig. S12A**). Sensory receptivity was determined using sound and luminance modulation indices (**Fig. 3B** and **C**, middle; **methods**), which identified sound-suppressed and sound-activated neurons (35.3%, **Fig. 3A** and **B**, top) concentrated near the tip of the probe (*i.e.*, in auditory areas, **Fig. 3D**), and luminance-suppressed or -activated neurons further up the shank in visual areas (26.9%, **Fig. 3C**, top and **Fig. 3D**). Decoding analyses confirmed that auditory, but not visual, units predicted sound stimulus presentation (**Fig. 3B**, bottom). In contrast, population vectors of visual neurons occupied distinct locations in a non-linear embedding (**Fig. 3A**, bottom) and could be used to reliably decode the luminance condition of different sessions, whereas A1 neurons could not (**Fig. 3C**, bottom). Seventy-five percent of auditory cortical neurons were modulated by behavior or white noise, of which 3-fold more were exclusively modulated by behavior (40%) than sound (12%), and 23% were tuned to both (**Fig. 3E**, left). A similar proportion was observed in visual cortices (though luminance was a coarser measure of sensory receptivity), with 42% of neurons tuned exclusively to behavior, 12% were luminance-sensitive, and 20% were tuned to both (**Fig. 3E**, right). We next compared the stability of behavioral tuning across light and dark conditions and found the features encoded in visual regions were more stable across light sessions than between light and dark sessions, with the most common tuning switches being to planar body motion and allocentric head movement in darkness (**fig. S12B**, left). Auditory neurons displayed greater stability across light-dark conditions, with the greatest changes occurring when unclassified and egocentric posture-responsive units acquired allocentric postural properties (**fig. S12C**, right).

**Fig. 3.**
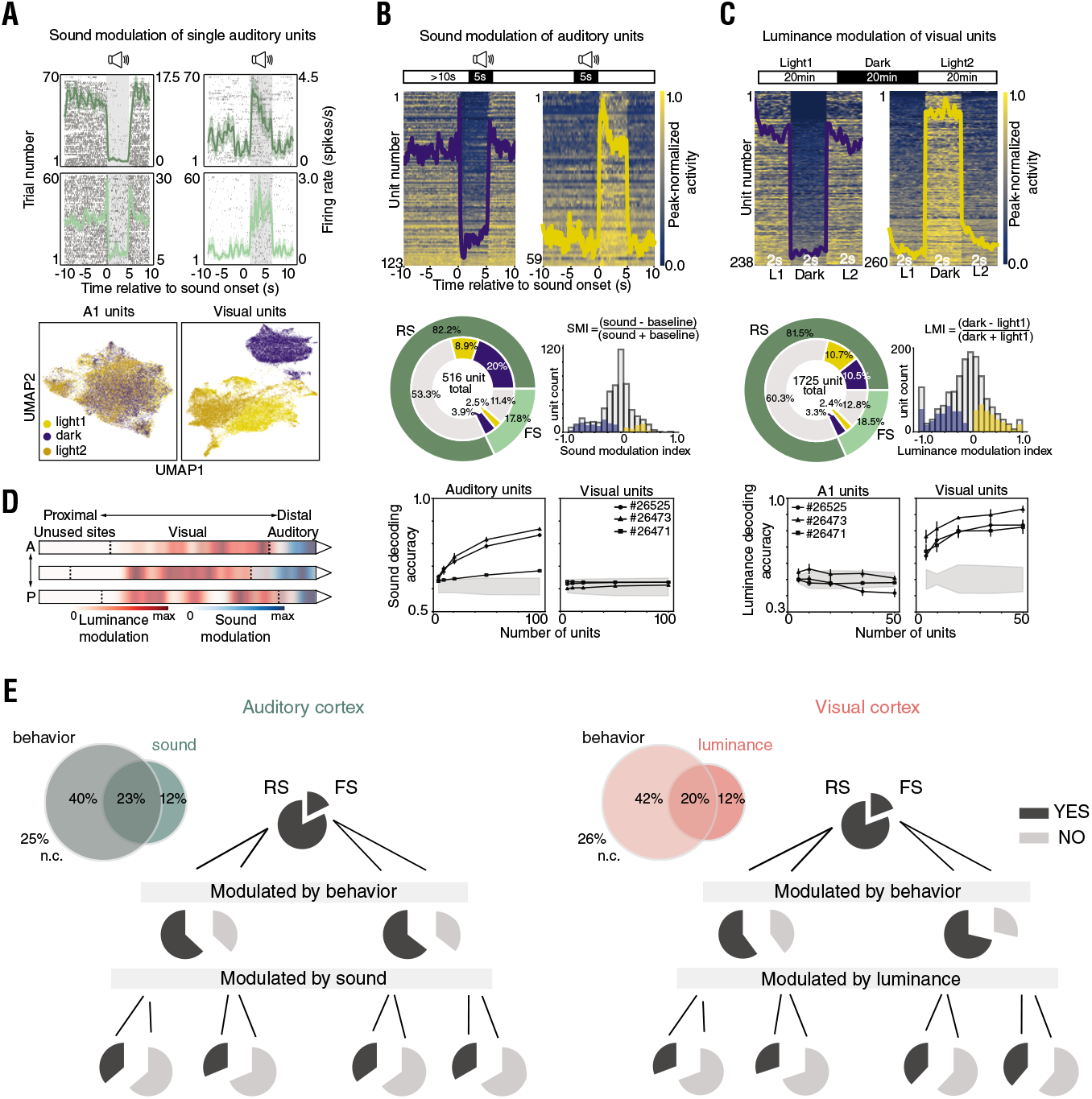
Prevalence and overlap of sensory and behavioral representation is similar across auditory and visual cortices. **(A)** (Top) Rasters and peri-event time histograms (PETHs) of sound-suppressed (left) and sound-excited (right) auditory cortical single units (green line and shading show trial averaged firing rate ±3 SEM; regular spiking (RS) units are in dark, and fast-spiking (FS) units in light green; sound stimulation in grey shading). (Bottom) Non-linear embedding of A1 population vector activity in an example rat (#26525); (right) same but for visual units. **(B)** (Top) White noise stimulation paradigm schematic. Trial averaged and ensemble averaged activity of all significantly sound-suppressed (left) and sound-excited (right) single units in auditory cortices during sequences of sound stimulation. (Middle left) Proportions of significantly sound-suppressed (dark blue) and sound-excited (gold) regular (dark green) and fast (light green) spiking auditory units. (Middle right) The sound modulation index distribution for all (grey), significantly suppressed (dark blue) and significantly excited (gold) auditory units. (Bottom) Decoding of sound stimulation with auditory (left) and visual (right) single units (symbols and vertical lines show mean decoding accuracy ±3 SEM for each rat; shaded area is 99% of the shuffled distribution). **(C)** (Top) Recording paradigm for light/dark sessions. 2 s segments of trial averaged and ensemble averaged activity of all significantly dark-suppressed and (left) dark-excited (right) single units in visual cortices. (Middle left) Proportions of significantly dark-suppressed (dark blue) and dark-excited (gold) regular (dark green) and fast (light green) spiking visual units. (Middle right) The luminance modulation index distribution for all (grey), significantly dark-suppressed (dark blue) and significantly dark-excited (gold) visual units. (Bottom) Decoding of luminance condition with A1 (left) and visual (right) single units (symbols and vertical lines show mean ±3 SEM of decoding accuracy for each rat, shaded area is 99% shuffled distribution). **(D)** Distribution of recording sites of single units modulated significantly by sound (blue) or luminance (red) across sensory cortices (opacity represents concentrations of units) in three sampled cortices. **(E)** (Left) Venn diagrams and pie charts summarizing the overlap between and breakdown of spiking profile, sound modulation and behavioral tuning (as determined by the GLM analysis; n.c. - non coding) in auditory cortices (A1 and A2D). (Right) Same as (left) but for luminance modulation and visual cortices (V1, V2L and V2M).

Recording large neuronal ensembles, usually several hundred units at a time, afforded the opportunity to seek putative synaptic connections and learn if characteristic connection types could suggest the role of behavioral modulation in each area. We found putative excitatory and inhibitory synaptic connections of varying strength in both auditory and visual (**Fig. 4A** and **B, methods**), and somatosensory and motor cortices (**fig. S13A** and **B**). We found different functional connection subtypes in each area (**Fig. 4C** and **fig. S13C-F**), but the most interesting results were in regions where sensory modulation was quantified. Relative to motor and somatosensory areas, visual pairs were more functionally homogeneous on average, and both visual and auditory synapses were weaker (**fig. S13E**). Furthermore, in visual areas, aside from excitatory (movement→movement, posture→posture) and inhibitory (movement⤏movement) communication between functionally homogeneous units, we found excitatory (posture→luminance modulated, posture→movement) and inhibitory (movement⤏posture) drive in functionally heterogeneous units (**Fig. 4C**, top). A notable majority of connections between heterogeneous units appeared in V2L (106/122 units, 86.8%), which was significantly larger than in V1 (*χ*^2^(1)=66.39, p=3.7e-16). Auditory areas, in contrast, exhibited completely different patterns of connectivity (**Fig. 4C**, bottom). On one hand, we found a significant amount of movement-inhibited posture-encoding units (*i.e.*, movement⤏posture), and this connection subtype occurred most prominently in A2D (14/18 units, 77.8%), as opposed to A1 (p=.015; one-sided binomial test). On the other, pairs of movement-inhibited sound-modulated units (*i.e.*, movement⤏sound modulated), featured more frequently in A1 (8/13 units, 61.5%), though the low total number of units precluded the difference with A2D to reach statistical significance (p=.29; one-sided binomial test). Finally, we noted that results obtained on synaptically connected pairs stood in stark contrast to those obtained on pairs receiving common input, which consisted almost exclusively of functionally homogeneous units (**fig. S13G**).

**Fig. 4.**
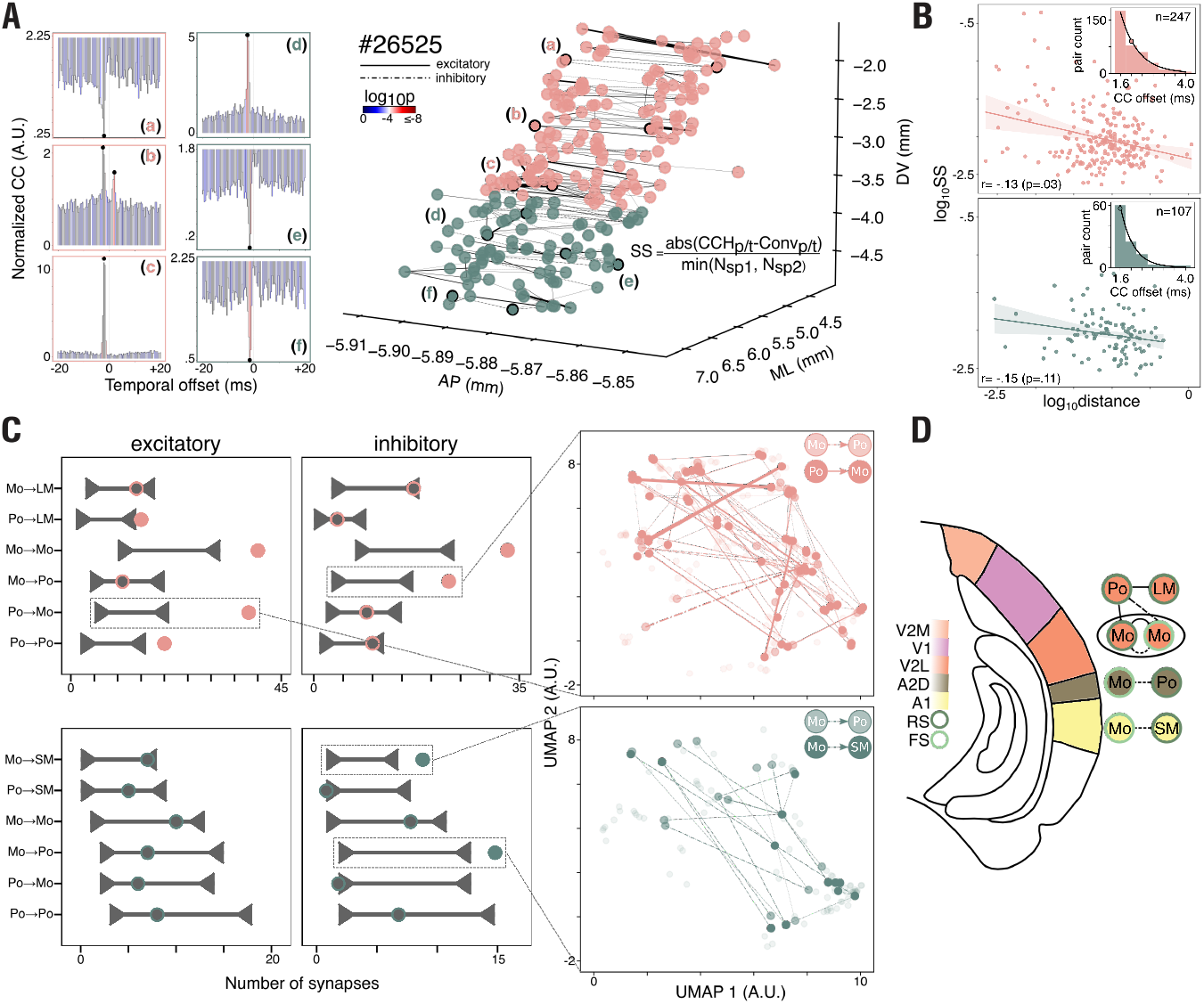
Synaptic connectivity patterns reveal behavioral information is employed differently across auditory and visual subregions. **(A)** (Left) Three example temporal spiking cross-correlograms from visual (a-c) and auditory cortices (d-f). (Right) All putative synaptic connections including outlined examples (a-f) from one example rat’s (#26525) visual (pink) and auditory (green) areas localized in anatomical space, with the width of each connection weighted by synaptic strength (SS). **(B)** (Top) The log-log relationship between anatomical distance and synaptic strength (SS) for all putative connections in visual cortices, (inset) distribution and median (colored circle) of cross-correlogram peaks/troughs. (Bottom) Same as above, only for auditory cortices. **(C)** (Left) Shuffled connection distributions (horizontal line with carets) and experimentally observed excitatory and inhibitory connections (circles) for various functional subtypes in visual (above) and auditory (below) cortices. (Right) Experimentally observed connections (outlined in left) in visual (pink) and auditory (green) cortices projected in a functional subspace (movement-responsive cells abbreviated to “Mo”; posture-responsive cells, “Po”; light modulated cells, “LM”; sound modulated cells, “SM”). We note that the outlined pair groups are largely separated. **(D)** (Top) The V2L network utilizes excitation and inhibition between postural and movement ensembles, potentially to coordinate impending movements with visual flow. (Middle) A2D fast spiking (FS) movement modulated ensembles inhibit gravity-relative, posture-responsive units, which could mediate sound localization. (Bottom) A1 FS ensembles inherit movement information, which could enable gain-modulation of local sound modulated regular spiking (RS) units in response to self-generated sounds.

## Discussion

In this study, we strove to address one of the core challenges in understanding ethologically relevant neural computations, namely, how cortical networks differentially represent freely-composed behavior. Visual and auditory areas were of particular interest because of their pervasive modulation by behavioral state (Niell and Stryker 2010; Reimer et al. 2014; Zhou et al. 2014; Osmanski and Wang 2015), movement during sensorimotor tasks (Keller et al. 2012; Ayaz et al. 2013; Polack et al. 2013; Saleem et al. 2013; Erisken et al. 2014; Fu et al. 2014; Schneider et al. 2014; Vinck et al. 2015; Williamson et al. 2015; Leinweber et al. 2017; Meyer et al. 2018; Vélez-Fort et al. 2018; Musall et al. 2019; Salkoff et al. 2019; Bouvier et al. 2020), or while animals cognitively idle in the dark (Stringer et al. 2019). To gain a clearer grasp on *which* actions drive cortical activity, we focused on what rats do when allowed to explore a familiar space without constraints. We converted tracked head and back points into series of postural and movement features, quantified the joint rat ethogram and characterized how recognizable modular actions were represented across cortical regions. The four overarching areas differed considerably in their connections, cytoarchitecture, and layers sampled, but our finding that every region carried sufficient information to decode nearly any action suggests that ongoing behavior continually modulates computations throughout dorsal cortical systems. This would serve to contextualize environmental inputs (Kaplan and Zimmer 2020) and inform internal predictive models (Friston 2005; Keller and Mrsic-Flogel 2018), since an animal’s behavior at any moment profoundly affects the spatiotemporal statistics of incoming sensory signals.

However, ethograms are descriptive of only one level of behavioral organization. When we “zoomed in” and modeled spiking responses on continuous elementary features, like rotational movements or angular head positions, we encountered a wealth of encoding variety across cortical structures. Compared to other areas, the response variance of primary motor cortex units was best captured by kinematic parameters (Georgopoulos et al. 1986; Pearce and Moran 2012), most conspicuously egocentric head and back movements and poses. The stability of the tuning curves for motor cortical units with or without added head weight was consistent with the spiking of the cells being linked more closely to the spatial position or movement path of the head, rather than generation of muscle forces to maintain or change head posture (Ward 1938; Crammond and Kalaska 1996; Kakei et al. 1999). Motor neurons most frequently encoded complex feature combinations, like head rotations around two or three axes, and had spiking activity accounted for better than any other area, as could be expected of a population generating sequences that control movement. In contrast, somatosensory neurons were best characterized by sparser models, usually back-related, which we trace to specific recording sites in the hindlimb region of S1 (Hall and Lindholm 1974; Halley et al. 2020), as back kinematics in quadrupeds are steadily affected by gross dynamics of the hindlimbs and hips.

Although world-centered encoding was the common denominator for visual and auditory cortices, the activity in visual ensembles was best described by allocentric horizontal motion of the head and body, which comports with studies demonstrating coding of angular head velocity (Vélez-Fort et al. 2018; Bouvier et al. 2020; Guitchounts et al. 2020b) and head direction (Guitchounts et al. 2020a) in rodent V1. In contrast, auditory neurons were most robustly triggered by gravity-relative head orientations. Such differences were made more prominent when we quantified connectivity differences between these areas and subregions (**Fig. 4D**). The lateral part of the secondary visual cortex (V2L), specialized for processing visual motion (Montero and Jian 1995; Andermann et al. 2011; Marshel et al. 2011), exhibited extensive cross-talk between movement-modulated cells, where putative feedforward excitation happened between similarly tuned cells, and apparent feedforward inhibition occurred between units encoding movements in opposite directions. More importantly, such fast spiking movement-responsive cells also inhibited posture-encoding units (*e.g.*, a left turn cell inhibiting a cell encoding rightward posture), and posture-modulated neurons were shown to excite luminance-modulated and movement-responsive cells (*e.g.*, a rightward pose cell exciting a leftward-rotation neuron). We suspect this could aid the ability to differentiate self-movements (McNaughton et al. 1994) from optic flow: if postural cells convey head stillness, their direct inhibition would indicate that visual scene changes are likely self-generated. Furthermore, when the animal’s head is fully extended in one direction, it can only really move in the opposite way; in this case, postural cells exciting opposite movement cells could aid in predicting the direction the visual scene will drift next (Nijhawan 2008; Hazoglou and Hylton 2019). However, determining the precise contribution of such circuit motifs during behavior will require future work manipulating functionally-identified neurons in controlled settings.

Lastly, the dissimilar connectivity patterns across primary and secondary auditory cortices may also indicate regional distinctions in how behavioral information is employed. Fast spiking cells encoding horizontal movements (*i.e.*, turning clockwise and counterclockwise) and inhibiting sound modulated units were mainly found in A1, which is strongly reminiscent of gain mechanisms for suppressing self-generated sounds identified in head-fixed animals (Schneider et al. 2014). Many such cells also showed evidence of receiving common input, consistent with efference copy circuits described in A1 (Reep et al. 1987; Budinger and Scheich 2009; Nelson et al. 2013), but here demonstrated in freely moving and sensing subjects. Movement-encoding fast spiking cells were also shown to inhibit allocentric posture-encoding cells, which mostly occurred in A2D (*e.g.*, a clockwise-turn cell inhibiting a cell encoding counterclockwise posture). We reason such a circuit could facilitate sound localization by strongly encoding minute changes in head position, which inform the rat about the position of the head and ears relative to the ground (Lauer et al. 2018). Gravity-relative head orientation may be of particular importance for animals with ear flaps, as such signals would convey the reliability of information arriving at each sensor (*i.e.,* ear), given their dissimilar concealment during natural motion.

In conclusion, detailed behavioral quantification will continue to reveal novel ethological insights (e.g. (Michaiel et al. 2020; Holmgren et al. 2021), but attention must be paid to lower level features as well, given the hierarchical and complex relationship of behavior and the brain. Defining how sensory cortices encode behavioral states and precise kinematics in freely moving animals opens the door to future investigations of how such networks respond in more naturalistically engaging environments, to establish how behavioral modulation is implemented in different circuits to solve locally relevant problems.

## Methods

### Subjects and electrode implantation

Experiments were performed in accordance with the Norwegian Animal Welfare Act and the European Convention for the Protection of Vertebrate Animals used for Experimental and Other Scientific Purposes. The study contained no randomization to experimental treatments and no blinding. Sample size (number of animals) was set *a priori* to at least three per recorded brain area, given the expected cell yield necessary to perform unbiased statistical analyses. A total of 7 male Long-Evans rats (age: 3-4 months, weight: 350-450 g) were used in this study. The rats were housed with their male littermates prior to surgery, and single housed in cages (45 × 44 × 30 cm) after surgery to avoid potential damage to the implants. All animals were kept on a reversed 12 h light-dark cycle and recordings were performed during their light cycle.

All 7 rats were implanted with silicon probes (Neuropixels version 1.0, IMEC, Belgium) developed for high-density extracellular recordings (Jun et al. 2017), targeting primary sensory and motor cortices. Each probe was coated with CM-DiI (Vybrant DiI, catalog #V22888, Thermo Fisher Scientific, USA) by repeatedly drawing a 2 μL droplet of CM-DiI solution at the tip of a micropipette up and down the length of the probe until the liquid dried, slightly changing the coloration of the shank. Subsequently, probes were, with electrical contacts facing up, stereotactically inserted into either the left hemisphere with a ~65°backward tilt in the sagittal plane to penetrate tissue from the posterior parietal cortex to primary motor cortex (n = 4, AP: −3.5 to −4.5 mm, ML: 1.9 to 2.7 mm) or, in the right hemisphere, with a ~50°lateral tilt in the coronal plane to target secondary and primary visual and auditory cortices (n = 3, AP: −5.52 to −6.5 mm, ML: 2.1 to 2.5 mm; see **fig. S1** for bregma-relative insertion coordinates of each probe). Probe insertions ranged from 3.9 to 7.2 mm in length across animals (**fig. S1**). External reference and ground wires were mechanically attached to a single skull screw (positioned at AP: +7 mm, ML: +2 mm) and sealed with a drop of SCP03B (*i.e.*, silver conductive paint). The remaining probe outside the brain was air sealed with a silicone elastomer (DOWSIL 3–4680 Silicone Gel Kit) and bead-sterilized Vaseline, and shielded by custom-designed black plastic housing to accommodate probes positioned at intended angles. Finally, the implant was statically secured with black-dyed dental cement to minimize light-induced electrical interference during recordings. Other steps of the surgical procedure are described in detail in our previous publication (Mimica et al. 2018). Following surgery, rats were subcutaneously administered fluids and postoperative analgesics and placed in a 37°C heated chamber to recover for 1-2 h prior to recordings.

### *In vivo* electrophysiology and behavior

Electrophysiological recordings were performed using Neuropixels 1.0 acquisition hardware, namely the National Instruments PXIe-1071 chassis and PXI-6133 I/O module for recording analog and digital inputs (Jun et al. 2017). Implanted probes were operationally connected via a headstage circuit board and interface cable above the head. Excess cable was counterbalanced with elastic string which allowed animals to move freely through the entire arena during recordings. Data were acquired with SpikeGLX software (SpikeGLX, Janelia Research Campus), with the amplifier gain for AP channels set to 500x, 250x for LFP channels, an external reference and AP filter cut at 300 Hz. In every session, signal was collected from all channels in the brain, typically from the most distal 384 recording sites (bank 0) first, followed by the next 384 recording sites (bank 1), consecutively.

Behavioral recordings were performed as individual rats foraged for food crumbs (chocolate cereal or vanilla cookies) scattered randomly into an octagonal, black open-field arena (2 × 2 × 0.8 m), with abundant visual orienting cues above and around the arena. All rats underwent a habituation phase prior to surgery during which they were placed on food restriction (to a minimum of 90% pre-deprivation body weight) to stimulate foraging behavior and were allowed to explore the arena daily. They were also acquainted and accustomed to the white noise presentations explained below. Food restriction was halted one day prior to surgery and recordings, by which time the animals were familiar with the environment. The entire data set for each animal was collected during 7-8 recording sessions (~20 min each) within the first 12 h (n=5) or 72 h (n=2) after recovery from surgery. The experiments were divided into two 4-session schedules in which recordings were made from bank 0 and bank 1, respectively. Each schedule consisted of the same ordering of conditions (light, dark, weight, and light/sound session). Each schedule started with a “light” session, where animals were run in dim lighting, followed by a “dark” session, in which all sources of light were either turned off or covered with fully opaque materials. Then, at the start of the “weight” session, a small copper weight (15 g) was attached to the animals’ implants before neural data was acquired. The last session of each schedule was either a “light” session or, when recording from auditory cortices, a “sound” session. During the latter, room lights were dimmed and white noise (5s duration) was played throughout the session at a pseudo-random inter stimulus interval (>10 s ISI) by a Teensy 4.0 Development Board controlled miniaturized Keyestudio SC8002B Audio Power Amplifier Speaker Module, running on custom-developed code. Between each schedule animals were returned to their home cage to rest.

### Perfusion and magnetic resonance imaging (MRI)

After recordings were completed rats received an overdose of Isoflurane and were perfused intracardially with saline and 4% paraformaldehyde. The probe shanks remained in the brains to give enhanced contrast and visibility during subsequent MRI acquisition. MRI scanning was performed on a 7T MRI with a 200 mm bore size (Biospec 70/20 Avance III, Bruker Biospin MRI, Ettlingen, Germany); an 86 mm diameter volume resonator was used for RF transmission, and a phased array rat head surface coil was used for reception. Brains were submerged in fluorinert (FC-77, 3M, USA) to remove background signal on the MRI. A 3D T1 weighted FLASH sequence was acquired at 0.06 mm^3^ resolution (TE: 10 ms, TR: 40 ms, NA: 4, matrix size: 360 × 256 × 180, FOV: 21.6 mm × 15.4 mm × 10.8 mm, acquisition time: 2 h 20 min).

### Histology and immunohistochemistry

After MRI scanning the shanks were carefully removed and brains were transferred to 2% dimethyl sulfoxide (DMSO, VWR, USA) for cryoprotection for 1-2 days prior to cryosectioning. All brains were frozen and sectioned coronally in 3 series of 40 μm on a freezing sliding microtome (Microm HM-430, Thermo Scientific, Waltham, MA). The first series was mounted directly onto Superfrost slides (Fisher Scientific, Göteborg, Sweden) and stained with Cresyl Violet. The second series was used to visualize Neuropixel tracks, labeled with CM-DiI, against neuronal nuclear antigen (NeuN) immunostaining, which provided ubiquitous labeling that enabled delineation of cortical layers. For immunostaining, tissue sections were incubated with primary anti-NeuN antibody (catalog no. ABN90P, Sigma-Aldrich, USA), followed by secondary antibody-staining with Alexa 647-tagged goat anti-guinea pig antibody (catalog no. A21450, Thermo Fisher Scientific, USA), after which the sections were rinsed, mounted, coverslipped and stored at 4°C. A more detailed immunostaining protocol is available per request. The third series of sections were collected and kept for long-term storage in vials containing 2% DMSO and 20% glycerol in phosphate buffer (PB) at −20°C. Using a digital scanner and scanning software (Carl Zeiss AS, Oslo, Norway), all brain sections were digitized using appropriate illumination wavelengths. The images were visualized with ZEN (blue edition) software and used subsequently along with MRI scans to locate recording probes in each brain.

### Probe placement

MRI scans were taken to locate the probes in 3D and to calculate the angle of each probe in the dorsal-ventral (DV) and medial-lateral (ML) axes. Since MRI scanning and histological staining were performed after perfusion with PFA, which cause a non-uniform reduction in brain volume, we reasoned the probe terminus would appear to have penetrated further in the tissue than was actually implanted. We therefore estimated the length of the implanted probe using the number of recording channels in the brain during the experiments. To locate the channel at which the probe exited the brain, we Fourier-transformed the median subtracted local field potential (LFP) signals at each channel along the probe and calculated power differentials between adjacent channels in the lower range frequencies (<10 Hz). We then located the largest shift in power between successive channels, which we identified as the point of exit from the brain (this analysis was adapted from the Allen Institute’s Modules for processing extracellular electrophysiology data from Neuropixels probes). The final length estimate for each probe was based on the identified surface channel and the physical geometry of the probe.

Probe placement was reconstructed in 3D (**fig. S1**) by first locating the entry point of the probe in the brain in CM-DiI-stained histological sections and their corresponding MRI scans. Given the probe length (calculated as elaborated above) and angles of the inserted probe relative to the tissue in different planes, we used trigonometry to calculate the rostral terminus of the probe (**fig. S1**). Anatomical coordinates were obtained by overlaying images of histological sections on corresponding sections from the rat brain atlas (Paxinos and Watson 2007). Probe tracks in the left and right hemisphere were followed from one coronal section to the next until the expected tip of the probe was reached, and area boundaries from the atlas were applied to determine the span of the probe in each brain region (grey line, **fig. S1D**). Using the within-region span and angle of each probe, we calculated the length, in micrometers, of each probe in each brain region in 3D, which allowed us to determine the number of channels in each region (with two channels spaced every 20 μm).

### Spike sorting and determining the spiking profile of single units

Given that the sessions were recorded in close temporal proximity, raw signals from recording files in each schedule (4 sessions) were concatenated in a unitary binary file, in order to keep the identity of each cluster across sessions. Spike sorting was performed with Kilosort 2.0 software using default parameters, followed by manual curation in Phy 2.0, where *noise* clusters were additionally separated from *good units* and *multiunit activity* based on inter-spike interval distributions, waveform features and the value of the Kilosort contamination parameter. Furthermore, good units were split into fast-spiking (FS) and regular-spiking (RS) subtypes by performing K-means clustering (where k=2) on spike width, peak-to-trough ratio, full width at half maximum and hyperpolarization (or end) slope data (**fig. S2**).

### 3D tracking, IMU and animal model assignment

The rats were tracked with seven retroreflective markers: four 9.5 mm spheres were affixed to a rigid body attached to the head (OptiTrack, catalog no. MKR095M3-10; Natural Point Inc., Corvallis, OR, USA), and three 9 mm circular cut outs of 3M retroreflective tape (OptiTrack, catalog no. MSC 1040) which were affixed to cleanly shaved locations on the trunk (Mimica et al. 2018). Their precise positioning was optimized to minimize interferences in picking up signals from individual markers. 3D marker positions were recorded at 120 fps with eight cameras (seven infrared and one B/W) 3D motion capture system (OptiTrack, Flex13 cameras & Motive v2.0 software). Additionally, a 9-DOF Absolute Orientation IMU Fusion Sensor (Adafruit, BNO055) was affixed to the implant chamber, such that angular velocities could be sampled directly and compared to tracking-derived features. The IMU data was acquired via custom-developed code through serial port terminal freeware (CoolTerm 1.7) at 100 Hz via another Teensy 4.0 Development Board, upsampled to 120 Hz *post hoc*, and rotated to match the reference axes defined by the tracking system. For precise alignment of acquired data streams, three additional infrared LED light sources were captured by the motion capture system. LED flashes (250 ms duration; random 250 ms ≤ IPI ≤ 1.5 s) were controlled by an Arduino Microcontroller C++ code which generated unique sequences of digital pulses transmitted to different acquisition systems throughout the recording and save the IPIs via serial port terminal freeware (CoolTerm 1.7). The detailed model assignment procedure has been described previously (Mimica et al. 2018). Briefly, all seven individual markers associated with the animal were labelled in a semi-supervised way using built-in functions in Motive. A rigid body was created using 4 markers on the head, and three markers on the body were labelled as separate markers. In addition to the markers on the animal, the three synchronizing LEDs were labeled as a separate rigid body (only marker sets with fixed distance over time can be labeled as a rigid body). After each session was fully labelled, remaining unlabeled markers were deleted and data were exported as a CSV file. The CSV file was converted to a format (pickle) compatible with our in-house graphical user interface (Mimica et al. 2018) for reconstruction of the coordinate system of the head from tracked points. Finally, tracking data was then merged with spiking data for further processing.

### Extracting postural variables from tracking data

Following the recordings, we labelled tracked points within the Motive (OptiTrack) interface, and imported labelled data into a custom script in Fiji. Using the four tracked points on the animal’s head, the geometry of the rigid body was estimated using the average pairwise distances between markers. We found the time point at which this geometry was closest to the average and used that time point as the template. We then assigned an XYZ coordinate system to the template with the origin located at the centroid of the four points, and constructed coordinate systems at each time point of the experiment by finding the optimal rigid body transformation of the template to the location of the head markers. To find the likely axis of rotation for the head (i.e. the base of the head), we found the translation of the coordinate system that minimized the Euclidean distance between the origin at time point t-20 and t+20, where t is measured in frames from the tracking system (120 fps). Next, the coordinate system was rotated to most closely match the Z-direction with the vertical direction of the room, and X-direction with that of the running direction, which was defined by horizontal movements of the origin from t-50 to t+50. Only time points where speed exceeded 10 cm/s were used to estimate running direction. The two objectives were combined by considering the sum of squared differences of the two sets of angles. This definition of running direction was used only to rotate the head direction, and was not used in subsequent analyses. Hyperparameters were chosen such that head placement using the resulting coordinate system visibly matched experiments.

To compute the postural variables for relating tracking to neural activity, we first denoted the allocentric angles of the head (pitch, azimuth and roll) relative to room coordinates, computed assuming the XYZ Euler angle method. We next denoted body direction as the vector from the marker above the root of the tail to the neck point. The egocentric angles of the head (pitch, azimuth and roll) relative to body direction were then computed assuming the XYZ Euler angle method. The back angles (pitch and azimuth) were determined relative to the horizontal component of body direction using standard 2D rotations, which were optimally rotated such that the average peak of occupancy was close to zero. The point on the neck was then used to determine neck elevation relative to the floor, as well as the horizontal position of the animal in the environment. Movement variables were estimated from the tracked angles using a central difference derivative with a time offset of 10 bins. Running speed was then estimated using a moving window of radius 250 ms. The values for self-motion were computed as the speed of the animal multiplied by the X and Y component of the difference in angles between the body direction at t-15 and t+15.

### Tuning curves to posture, movement and navigational variables

Angular behavioral variables were binned in 5°, with the exception of back angles, which were lowered to 2.5°. Movement variables were binned in 36 equally-spaced bins, spanning the range of recorded variables such that there was a minimum occupancy of 400 ms in both the first and last bins. Neck elevation bins were 1 cm, while position in the environment was estimated using 6.67 cm bins. Finally, self-motion used a bin size of 3 cm/s. For all rate maps, the average firing rate (spk/s) per bin was calculated as the total number of spikes per bin, divided by total time spent in the bin. All smoothed rate maps were constructed with a Gaussian filter with a standard deviations of 1 bin. Only bins with a minimum occupancy of 400 ms were used for subsequent analysis. To shuffled receptive field distributions, we shifted the neural activity 1000 times on the interval of ±[15,60] s and recomputed tuning curves for each variable.

### Defining composite actions

The behavioral clustering pipeline is sketched in **Fig. 1B** and adapted from prior work (Berman et al. 2014). The starting point is the time series of postural parameters and the running speed of the animal, **Fig. 1B** (panel 1). Running speed, neck elevation and back Euler angles (pitch and azimuth) were defined as explained above (see “Extracting postural variables from tracking data”), while 3D head direction relative to the body direction was parameterized using the exponential map (Grassia 1998). The time series of each of the 7 variables (6 postural parameters plus running speed) was detrended using 3^rd^ degree splines with equally spaced knots at 0.5 Hz, **Fig. 1B** (panel 2). Time frequency analysis was then performed on the detrended time-series for each of the original variables using Morlet wavelets, at 18 Fourier frequencies dyadically spaced between 0.5 and 20 Hz, **Fig. 1B** (panel 3). The square root of the power spectral density was centered and rescaled dividing it by the variance of the smoothed signal (fit resulting from the spline interpolation). The smoothed signal was z-scored and concatenated with the rescaled spectrogram yielding a 133D feature vector for each tracked time point (120 Hz). Feature vectors were downsampled in time at 1 Hz and pooled across animals and conditions. Principal Component Analysis (PCA) was performed, indicating that the first 22 principal components explained 97.2% of the variance. Only these 22 principal *eigenmodes* were retained and the dimensionality was further reduced to 2 via tSNE (Maaten and Hinton 2008) embedding (Euclidean metric, perplexity=200), **Fig. 1B** (panel 4). The embedding was then used to estimate a probability mass function (PMF) on a 60×60 lattice in the 2 tSNE dimensions by convolving the raw histogram with a two-dimensional Gaussian (width=1.). We segmented the 60×60 lattice by applying a watershed transform (Meyer 1994) to the additive inverse of the PMF, **Fig. 1C**. All data points falling within a watershed-identified region were assigned the same action label. Timepoints not belonging to the dataset used for PCA were classified by minimizing the Euclidean distance of the feature vector in the 22-dimensional PC space from the datapoints used for training. Names were attributed to the action labels after post-hoc visual inspection of individual timestamps in one session per animal in a graphical user interface (GUI) and by comparison to the postural decomposition of the behavior (**fig. S4A** and **B**; **movies S1-6**).

### Encoding and decoding of actions

The average firing rate (spk/s) of each cell per attributed label was calculated as the total number of spikes emitted during the action, divided by total time spent in it. Additionally, the average firing rate of the cell was computed separately in two halves of the dataset, the two halves constructed to include half of the timepoints spent in each action. To compare with shuffled data, we shifted the neural activity 1000 times on an interval of ±[15,60] s. Shuffled distributions were also constructed for each of the two halves of the dataset. A cell was classified as encoding a behavior if these 2 criteria were met: [i] its average firing rate at the behavior on the whole dataset was either (a) below the 0.01^th^ percentile, or (b) above the 99.99^th^ percentile of the shuffle distribution [ii] if (a), its average firing rate for the action in both halves of the dataset was below the 2.5^th^ percentile of the shuffle distribution of each half respectively, while if (b), its average firing rate for the behavior in both halves of the dataset was above the 97.5^th^ percentile of the shuffle distribution of each half respectively. For [i] the 99% significance level was Bonferroni-corrected for multiple comparison.

Spike counts time series were constructed by counting the number of spikes fired by a cell in each 8.33 ms time bin. Behavioral decoding from spike count data was performed on every session for which more than 10 cells were simultaneously recorded. A naïve Bayes classifier was trained on all the actions with an occupancy larger than 16 s, resulting in 34 ± 3 (mean ± SEM) actions per session to be decoded. Decoding was performed on 20 samples of 400 ms each (50 bins) per action, while the rest of the dataset was used for training. The classifier consisted of a binomial likelihood and a categorical prior determined by the occupancy of actions in two different randomly selected sessions of the same animal. We defined the decoding accuracy for an action as the fraction of samples whose label was correctly classified. For comparison we also classified actions using only the prior distribution.

### GLM and model selection

We binned the spike train of all neurons with 8.33 ms time bins to match the tracking frequency of 120 Hz. Let *y*_*t*_, *t* = 1, … , *T* be the binarized spike count of a neuron in time bin *t* of a total of *T* in the whole recording session; *y*_*t*_ = 1 indicates that the neuron emits one or more spikes in bin *t*, whereas *y*_*t*_ = 0 indicates that the neuron doesn’t fire in bin *t*. The probability of *y*_*t*_ is given by a Bernoulli distribution,

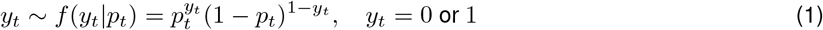

where *p*_*t*_ is the probability that *y*_*t*_ = 1. Let χ_*t*_ = {*x*_1_(*t*), … , *x*_*m*_(*t*)} represents the *m* tracked and factorised features at time t: nine postural features (pitch, azimuth and roll of the head in allocentric and egocentric reference frames, back pitch and azimuth, neck elevation), their first derivative values, body direction, speed, position and self-motion. For each feature *i*, let *x*_*i*_(*t*) be a binary vector of length *N*_*i*_ (number of bins used to factorize covariate *i*: 15 for 1D features; bin size of 5 cm/s for self-motion and 10 cm for position), whose components are all 0, but the one corresponding to the bin in which the features falls at time *t*. To study how well a neuron can be explained by one or more features, we fit the activities of a single neuron using generalized linear models *M* (Nelder and Wedderburn 1972) with logit link function,

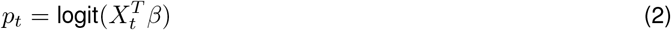

where *β* are the parameters of the model and 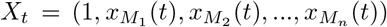 is the vector of *n* features included in model *M*. Thus the log-likelihood of the model is,

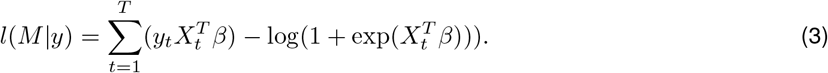

We estimated all models with an additional L1 regularization with the learning rate *λ*=10^−4^.

To determine which subset of these features best explain the neural activity, we performed a forward selection procedure (Hardcastle et al. (2017) combined with a 10-fold cross-validation scheme. For each neuron, we first partition the data into 10 approximately equally sized blocks *a* = 1, .., 10, where each block {*y*^*a*^, *X*^*a*^} consists of consecutive time bins. We then computed the average held-out scores across folds. The initial simple model consisted only of an intercept and features were added sequentially through three-steps:

1. For each feature not included in the model, and each fold *a*, we fitted the GLM with the feature added, *M*, on 9 data blocks and computed the log-likelihood *l*(*M* |*y*^*a*^) for the test data. After iterating over folds we took the average over folds of 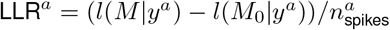, where 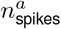 is the number of spikes in the test data of fold *a*, while *l*(*M*_0_|*y*^*a*^) is the out-of-sample log-likelihood of the intercept model in fold *a*. We determined which feature had the largest value of the average LLR and selected it as a candidate feature to be included in the model.
2. For the given candidate feature, we employed a one-sided Wilcoxon signed rank test on the out-of-sample log-likelihood across folds *l*(*M*|*y*^*a*^) of the more complex model and the current model. The null hypothesis is that the more complex model yields smaller or equal values of *l*(*β*|*y*^*a*^) with respect to the less complex model.
3. If the null hypothesis was rejected (*α* = .01), the new feature was added to the model and the forward selection procedure continued. The selection process stopped when the null hypothesis was not rejected or no more features were available.

After a final model is selected for each cell, we calculated the contribution of each feature belonging to the final model via 10-fold cross-validation. Assuming that the final model *M*_full_ includes n covariates, for each selected covariate *x*_*i*_, *i* = 1, … , *n*, we considered the partial model 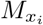 which includes all the covariates except *x*_*i*_, and the intercept model *M*_0_. Then for each partition *a* of the data we trained the three models and computed the out-of-sample log-likelihoods 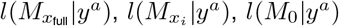. Finally we define the contribution of each covariate to the final model as the relative log-likelihood ratio

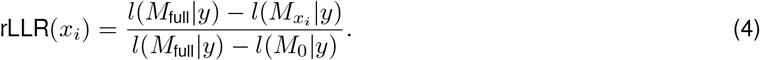

where *l*(*M*|*y*) is the average across folds *a* of *l*(*M*|*y*^*a*^).

We measure the prediction accuracy of a model *M* relative to the intercept model *M*_0_ as the average across folds *a* of McFadden’s pseudo-R^2^ (McFadden 1973)

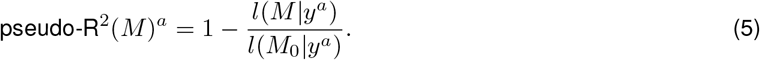

### Anatomical topography of tuning features

Data from the three left hemisphere-implanted animals were used to compute anatomical gradients of behavioral tuning across adjacent brain regions (the fourth rat, #26148, was excluded due to limited anatomical coverage). Since the exact anterior-posterior (AP) placement of the probes differed across animals, the overall extent of the three probes was calculated using the most posterior and the most anterior anatomical locations. This physical distance comprised the rows of a matrix with 3 columns, in which each column represented data from individual animals. Based on where the probe was located in this physical space and how many cells of a particular tuning type (*e.g.*, allocentric head roll) were recorded, multiple matrices were created for each feature. These matrices were then divided into 7 equal segments, each corresponding to 1 mm of tissue, and the numbers of cells tuned to particular features were summed. Since the absolute number of cells varied across animals, the data were presented as a proportion of the total number of cells recorded in a given 1 mm segment. We applied a *χ*^2^ test to determine if the observed distribution of cells tuned to each feature was significantly different from a uniform distribution over the cortical surface. The features with the most significant differences were plotted in **fig. S9A**.

All three right hemisphere-implanted animals were used for this analysis. Instead of calculating the absolute spatial extent of the three probes along the medial - lateral (ML) axis, the probes were aligned based on the auditory/visual border. The same approach was applied as above, and the distribution of the proportion of cells tuned to each feature over the anatomical surface was calculated, and significance was assessed using the *χ^2^* test. The features showing the strongest anatomical differences were plotted in **fig. S9B**.

### Sensory modulation indices and decoding

To obtain peri-event time histograms (PETHs) relative to sound stimuli onset, each spike train was zeroed to tracking start, purged of spikes that exceed tracking boundaries and binned to match the tracking resolution. It was further resampled (to 50 ms bins) to encompass a large window (10 s) before and after every event onset (the start of the white noise stimulation). Spike counts were converted to firing rates (spk/s) and smoothed with a Gaussian kernel (sd=3 bins). To identify sound responsive units, we calculated the sound modulation index (SMI) for each cell on PETHs averaged across all trials (**Fig. 3B**, top and middle). SMI is the difference between the “sound” (500-1000 ms post-stimulation) and the “baseline” firing rate (1000-500 ms pre-stimulation) divided by the sum of the two, such that a negative SMI signifies higher firing before, and a positive SMI signifies higher firing following sound onset. The statistical significance of each SMI was determined with a Wilcoxon signed-rank test (p<.05) performed on all “sound” and “baseline” trial sequences.

We used a nearest neighbor decoder to query whether we could predict the sound being “on” or “off” given only auditory or visual ensemble activity, for each rat separately (**Fig. 3B**, bottom). The sound event vector (“on” or “off”), together with the spike train of each single cell, was resampled to 10 Hz resolution and the latter were convolved with a Gaussian kernel (sd=1 bin). In each run (for a total of 100 runs per unit number) we pseudorandomly subsampled either 5, 10, 20, 50 or 100 different cells and divided the data into 3 folds, where each third of the data once served as the test and the other two thirds as the training set. We calculated Pearson correlations between every test set ensemble activity vector and every ensemble vector in the training set. This enabled us to obtain a predicted sound stimulus value (“on” or “off”) for each test frame by assigning it the sound stimulus value of the highest correlated training set vector. Decoding accuracy was defined as the proportion of correctly matched stimulus states across the entire recording session (theoretically varying from 0 to 1). To obtain the null-distribution of decoded accuracy we shuffled the spike trains of each subsample in the first run 1000 times (as described above) and followed the same described procedure that resulted in shuffled accuracy distributions.

Since “light” and “dark” conditions were not varied on a trial, but on a session basis, we computed PETHs by first searching for all ≤2 s time windows where the speed of the animal was ≤5 cm/s, effectively equating to quiescence or epochs of slow movement. We did this in three sessions: light1, dark and light2 by subsampling the number of events from the session the had the fewest such events in the other two sessions. The firing rates (spk/s) in each window bin were calculated using the same method described above. To identify luminance responsive units, we calculated the luminance modulation index (LMI) for each cell on PETHs averaged across all trials. (**Fig. 3C**, middle). LMI is the difference between the “dark” (full 2 s window) and the “light1” firing rate (full 2 s window) divided by the sum of the two, such that a negative LMI signifies higher firing in light conditions, and a positive LMI signifies higher firing in the dark condition. The statistical significance of each LMI was determined with a Wilcoxon signed-rank test (p≤.05) performed on all “dark” and “light1” trial sequences, provided that the same test yielded no difference in firing rates (p>.05) between “light1” and “light2” conditions. To visualize these differences, we concatenated the population vectors (all recorded cells in A1 or V) of all three sessions (in one example animal), z-scored them, and performed the principal component analysis (PCA). We determined the vertex (or the “knee”) of the scree plot and selected all components preceding it for non-linear low-dimensional embedding of individual timepoints with UMAP (**Fig. 3A**, bottom).

To determine the relative strength of the sound and luminance modulation along the recording probe, we counted all significantly modulated units (both suppressed and excited) at their respective peak channels, joined all channels of 2 successive rows in one count (totaling 4 channels every 2 rows), and normalized this count by the maximal count obtained.

The nearest neighbor decoding was also adjusted to accommodate for the lack of a trial based structure. First, due to the fact that the secondary auditory area (A2D) had multisensory properties, i.e., units which were sensitive to both sound and luminance change, we focused our analysis on primary auditory (A1), together with all recorded visual (V1, V2L and V2M) neurons. The data were downsampled and smoothed as described above, and synthesized by taking the last quarter of timepoints in the light1 session, the temporally adjacent first quarter of timepoints in the dark session and the temporally distant second half of the light2 session. Similarly to the sound decoder, in each run (for a total of 100 runs per unit number) we pseudorandomly subsampled either 5, 10, 20, 35 or 50 different cells (adjusting for the lower total number of cells in A1). For each test set ensemble activity vector (i.e., the second half of light2) we computed Pearson correlations to every ensemble vector in the training set (light1 + dark) and obtained a predicted condition status by assigning it the condition status of the most highly correlated training ensemble vector. Since luminance did not change within a session, shuffling spike trains would not suffice (because it would not eliminate the overall lower/higher rate relative to the other session), we randomly permuted the ensemble vectors across the training set (i.e., light1 + dark) at each time point 1000 times in the first run which resulted in the null-distribution of decoded accuracy.

### Weight and behavioral tuning

To ascertain whether weight had a behavioral effect on the measured variables, we primarily focused on head-related features, neck elevation and speed, assuming these would be affected the most. For each feature we computed differences between the total occupancy in every bin between the weight and light2 sessions, across all rats and looked whether the 99% CI of these differences overlapped with zero.

To estimate whether adding the weight on the head had any affect on the neural coding or activity, we performed several analyses. Since our recordings were performed over multiple successive sessions, we analyzed the change in the overall activity of spiking through time. Therefore, in each cluster, spikes are allocated to broad 10 s bins and smoothed with a Gaussian kernel (sd=1 bin). They were then concatenated into a single array and a rolling mean (size=50 bins) was calculated over the whole window for display purposes. The “baseline” firing rate was defined as the total spike count within a session divided by the total session time. A “stable” baseline rate was the weight session rate above .1 spk/s whose difference to the reference session rate (light2 sessions were picked as reference sessions as overall rates tended to be more similar to the weight sessions) was smaller than 20% of its own rate. To obtain the shuffled distribution of rate differences, we pseudorandomly permuted individual cells’ rate identities across light1, weight and light2 sessions 1000 times. Our subsequent analyses focused on the effect of weight on different tuning features, namely the observed differences in areas under the tuning curve (AUC), the observed differences in information rate (Skaggs et al. 1993), the observed differences in the stability of tuning curves, and the observed differences in tuning peak positions. To determine whether any of these difference distributions were significantly different compared to a null-distribution, we created shuffled distributions of differences by pseudorandomly permuting session identities of the data 1000 times and recomputing the differences.

### Functional connectivity

Spike trains were binned in .4 ms wide bins and dot products (cross-correlograms, CCG) were computed between every spike array and any other jointly recorded spike array with temporal offsets spanning the [-20, 20] ms range with .4 ms steps. To generate a low frequency baseline cross-correlation histogram for comparison, the observed CCG was convolved with a “partially hollowed” Gaussian kernel (Stark and Abeles 2009), with a standard deviation of 10 ms, and a hollow fraction of 60%. The observed coincidence count (CCG) is compared to the expected one (low frequency baseline) which is estimated using a Poisson distribution with a continuity correction, as previously described (English et al. 2017). A putative connection was considered synaptic if the following conditions were met: 99.9999/0.0001 (for excitatory/inhibitory connections, respectively) percentile of the cumulative Poisson distribution (at the predicted rate) was used as the statistical threshold for significant detection of outliers from baseline, (2) two consecutive bins needed to pass the threshold within the ±1.6-4 ms window (Senzai et al. 2019), and (3) there should be no threshold passing in the ±1.6-0 ms range. Alternatively, if the peak/trough occurred in the ±1.6-0 ms range, and two consecutive bins passed the threshold for detecting outliers, the units were considered as receiving common input.

The 3D position of each neuron in a connected pair was determined by first computing its center of mass on the probe surface, based on peak absolute template waveform amplitudes on the peak waveform channel and 20 adjacent channels below and above the peak. The exact DV, ML and AP positions were then computed taking into account the insertion site, the total length of the probe in the brain and its angles in the tissue, as explained above. Synaptic strength was defined as the absolute difference between the spike coincidence count at the CCG peak/trough (for excitatory/inhibitory connections, respectively) and the slow baseline at peak/trough, normalized by the minimum number of spikes between the two spike trains (*i.e.*, the theoretical maximum number of coincidences).

All neurons in the dataset were assigned with a variable that best fit its spiking variability in the dark session, based on the mean cross-validated relative log-likelihood ratio (rLLR) of single covariate models relative to the null model (23 behavioral features + unclassified cells). The “functional space” map was obtained for visualization purposes only by performing PCA on a matrix containing such values for all 23 covariates, in addition to the SMI and LMI estimates and p-values. The first n components that cumulatively accounted for 90% of the variance were then embedded on a 2D plane by uniform manifold approximation and projection (UMAP). The 24 feature list was further simplified by grouping variables in 11 categories: unclassified, position, speed-related, egocentric head posture, egocentric head movement, allocentric head posture, allocentric head movement, back posture, back movement, neck elevation, neck movement. Variables were plotted on log-log scales for visualization purposes only, but all presented statistics (Mann-Whitney U test was chosen, as Levene’s test established groups had unequal variances) were performed on the original data, and “functional distance” was calculated across the original 28 variables (23 covariates + SM and LM indices and p-values).

Excitatory and inhibitory connection pairs were classified in 6 broad functional categories: (1) movement modulated neuron preceding a sensory modulated cell (either sound or luminance, for auditory and visual ensembles, respectively), posture modulated neuron preceding a sensory modulated cell, (3) movement modulated neuron preceding a movement modulated cell, (4) movement modulated neuron preceding a posture modulated cell, (5) posture modulated neuron preceding a movement modulated cell, and (6) posture modulated neuron preceding a posture modulated cell. Assessment of whether the connection pair numbers in each category could have been observed by chance was done by subsampling pseudorandomly paired units 1000 times, provided that: (1) the anatomical distance between cells was shorter or equal to the maximal one observed in the true data, (2) there were equal numbers of excitatory and inhibitory connections as in the real data in each run, (3) the connection was physiologically plausible (excitatory/inhibitory connections could only be formed in the RS/FS cell was the presynaptic neuron, respectively). Lastly, we statistically tested whether the observed difference in neuron counts between subregion categories (either A1-A2D, or V1-V2L) was larger than theoretically expected by an equal split (50%-50%), using the *χ*^2^ goodness-of-fit test if the expected frequencies exceeded 5 in each category, or the binomial test otherwise.

### Data and code availability

All datasets generated for this project, together with supporting documentation, are available for download at <u>fig**share**</u>. The code pertaining to the experimental pipeline for data acquisition and preprocessing can be found at **https://github.com/bartulem/KISN-PyLab**. The code used to analyze the data and make the figures can be found at **https://github.com/bartulem/KISN-pancortical-kinematics**.

## Author contributions

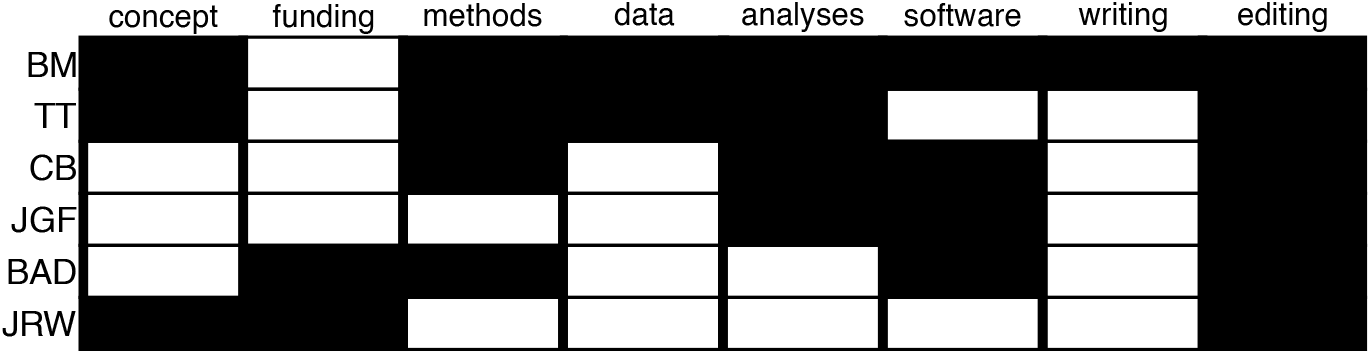

## Acknowledgements

We thank D. Deutsch, A.L. Falkner, M. Long and M. Murthy for helpful comments on the manuscript; M. Andresen for tissue sectioning; R. Gardner, T. Waaga and V. Normand for the initial assistance with Neuropixels recordings; W. Zong for assistance in designing the 3D printed cap housing the Neuropixels probes; S. Gonzalo Cogno, R. Gardner, V. Normand and T. Waage for fruitful discussions along the way; K. Haugen and H. Waade for technical and IT assistance; M. Witter and M. Nigro for assistance with anatomical delineations; D. Hill and M. Widerøe of the Norwegian University of Science and Technology (NTNU) MRI Core Facility and S. Eggen for veterinary oversight. This project used the high performance computing infrastructure, IDUN, provided through the NTNU (Själander et al. 2021). We also wish to express our gratitude towards individuals, teams, organizations and funding programs behind fundamental packages for scientific computing and visualizations in Python which relieved our workload substantially: Matplotlib (Caswell et al. 2021), Numba (Lam et al. 2015), NumPy (Harris et al. 2020), Pandas (Reback et al. 2021), Scikit-Learn (Pedregosa et al. 2012), and SciPy (Virtanen et al. 2020).

---

**Fig. S1.**
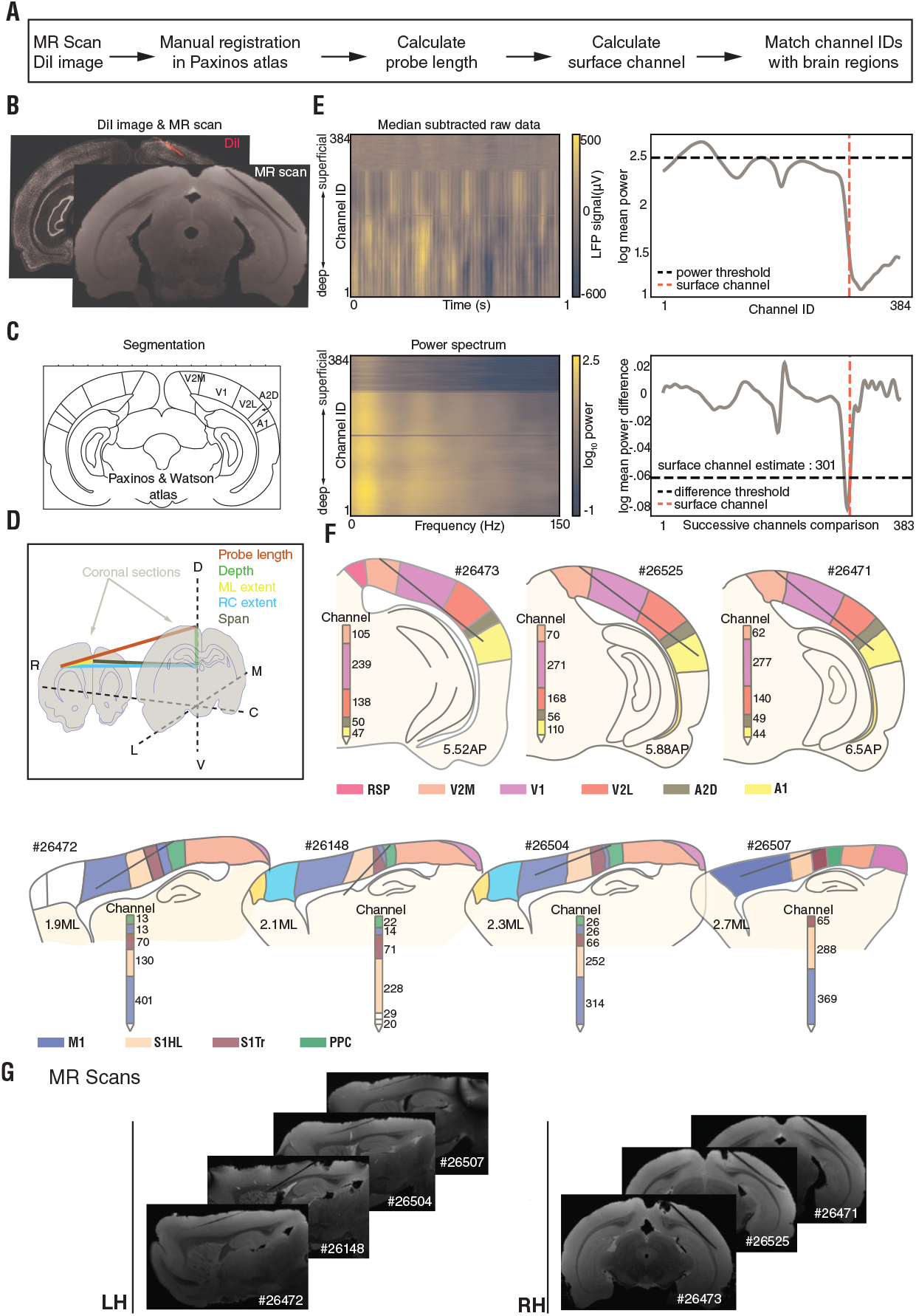
Probe and channel localization in sensory and motor cortices. **(A)** Pipeline for localizing Neuropixels 1.0 probes and recording sites in specific brain regions. **(B)** Coronal section of a CM-DiI and NeuN co-stained section (back) and MR-scan of the same brain prior to sectioning (front) with a probe spanning visual and auditory cortices. **(C)** Atlas images (Paxinos and Watson 2007) were used to determine regional boundaries for coronal sections containing CM-DiI-stained probe tracks. **(D)** Schematic showing how probe placement (red) was registered in 3D space with respect to dorsoventral (DV; green), anteroposterior (AP; blue), and mediolateral (ML; yellow) axes. **(E)** The surface channel was located using electrophysiological measures across all recording sites. (Upper left) The mean subtracted LFP signal and (lower left) power spectrum across recording channels change abruptly where the probe exits the brain (1 s of data is shown). (Upper right) Power fluctuations across low band frequencies (<10 Hz) with an arbitrary threshold (black dashed line) and surface channel estimate (red) obtained from power differentials, below. (Lower right) Differences in power between successive channels with a log_10_ mean power difference threshold no higher than −.06 (black) indicate the surface channel as 301 (red). **(F)** Anatomical reconstructions of probe placements based on probe length and channel count estimates. (Top row) Probe track reconstructions in the coronal sections from the brains of three rats with probes spanning primary and secondary visual and auditory areas in the right hemisphere; sections are arranged from anterior (left) to posterior (right). (Bottom row) Sagittal reconstructions of probe tracks in the left hemisphere of four rats; sections are arranged from medial (left) to lateral (right). Dark gray lines denote the probe location in each brain, and the number of channels in each region are shown on schematic probes inset with each reconstruction. **(G)** MR scans showing the probe locations in each of the seven animals. LH shows probe placement in primary somatosensory and motor cortices in the sagittal plane, whereas RH shows probe placement in visual and auditory regions in the coronal plane.

**Fig. S2.**
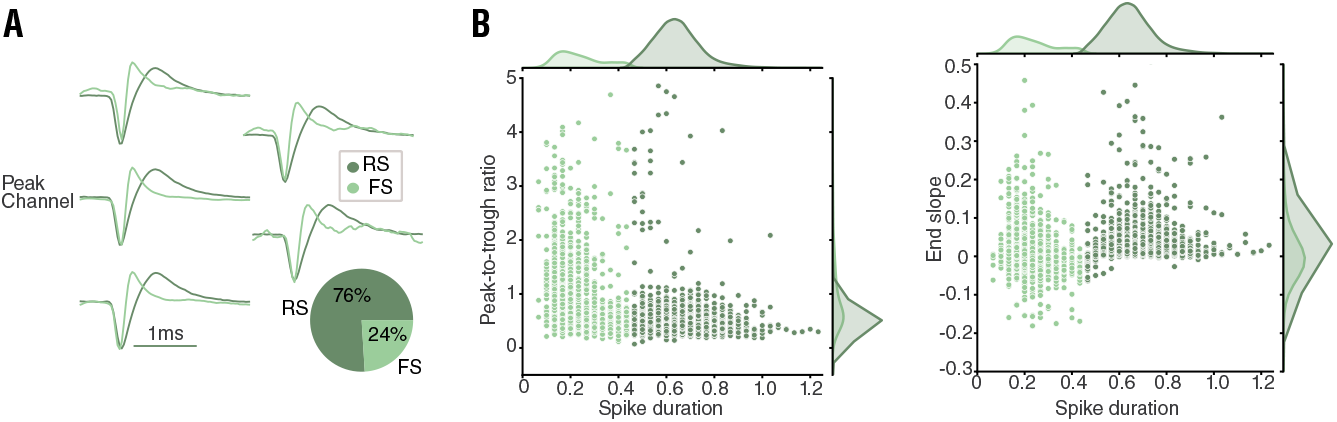
Classifying single units on the basis of spiking profiles. **(A)** Mean waveforms of two example clusters on the same set of adjacent channels: fast spiking (FS) cluster in light, and regular spiking (RS) cluster in dark green; (bottom right) overall breakdown of FS and RS clusters in the entire dataset. **(B)** (Left) FS and RS peak-to-trough ratio and spike duration distributions. (Right) FS and RS end slope and spike duration distributions.

**Fig. S3.**
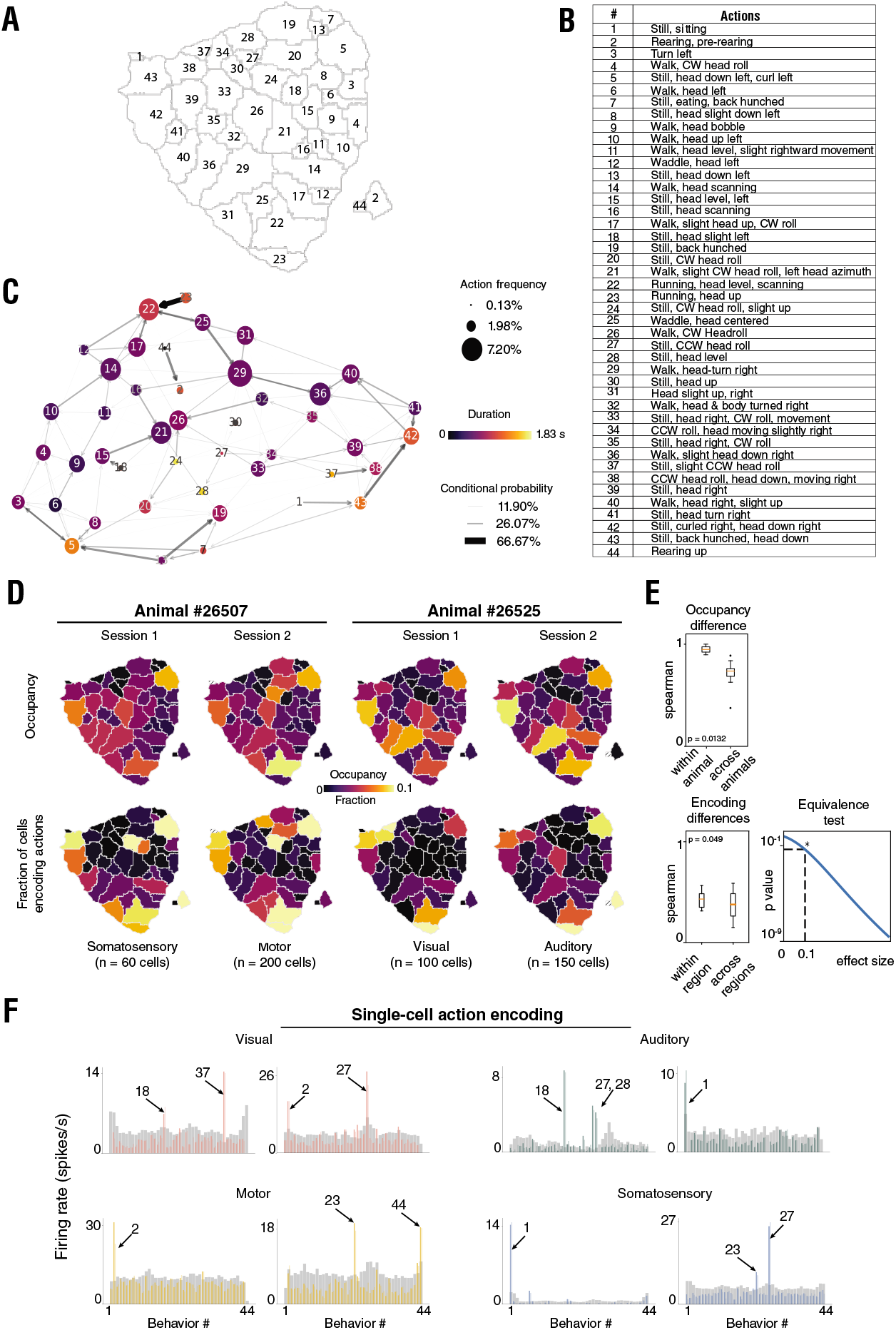
Statistics of actions and variability of encoding of actions across cortical areas. **(A)** Segmented t-SNE behavioral space with numerical labels and **(B)** description of each action. **(C)** The frequency of each action is shown via node size; color indicates mean action duration (0.49±0.36 s, mean ± SD across actions); time-conditional probability of the subsequent actions is indicated by arrow transparency and thickness. **(D)** (Top) Relative occupancy of actions in the segmented t-SNE behavioral space from two example animals. (Bottom) Fraction of cells encoding each action (see **methods**). **(E)** (Top) Spearman’s *ρ* of relative occupancy of actions between sessions of the same animal 0.94±0.06 (mean ± SD; n=10; box plots extend from the 25^th^ to 75^th^ percentile of the sample distribution; orange lines indicate the median; whiskers span the data range; outliers (black dots) were excluded), and between sessions of different animals 0.71±0.13 (mean ± SD; n=68; p=.0132, Mann-Whitney U test for difference between medians in the two groups). (Bottom left) Spearman’s *ρ* of the fraction of cells encoding each action between recordings from the same brain region 0.40±0.14 (mean ± SD; n=38; p=.004, permutation test), and between recordings from different brain regions 0.36±0.16 (mean ± SD; n=98; p=.008, permutation test). (Bottom right) p-value of two one-sided t-test statistics for the equivalence between means of the Spearman’s *ρ* in the two groups as a function of effect size (*Cohen’s d*; *: p=.049 at *Cohen’s d* effect size=0.1). **(F)** Examples of single-cell behavioral encoding in each of the four overarching cortical regions, with action identity indicated above.

**Fig. S4.**
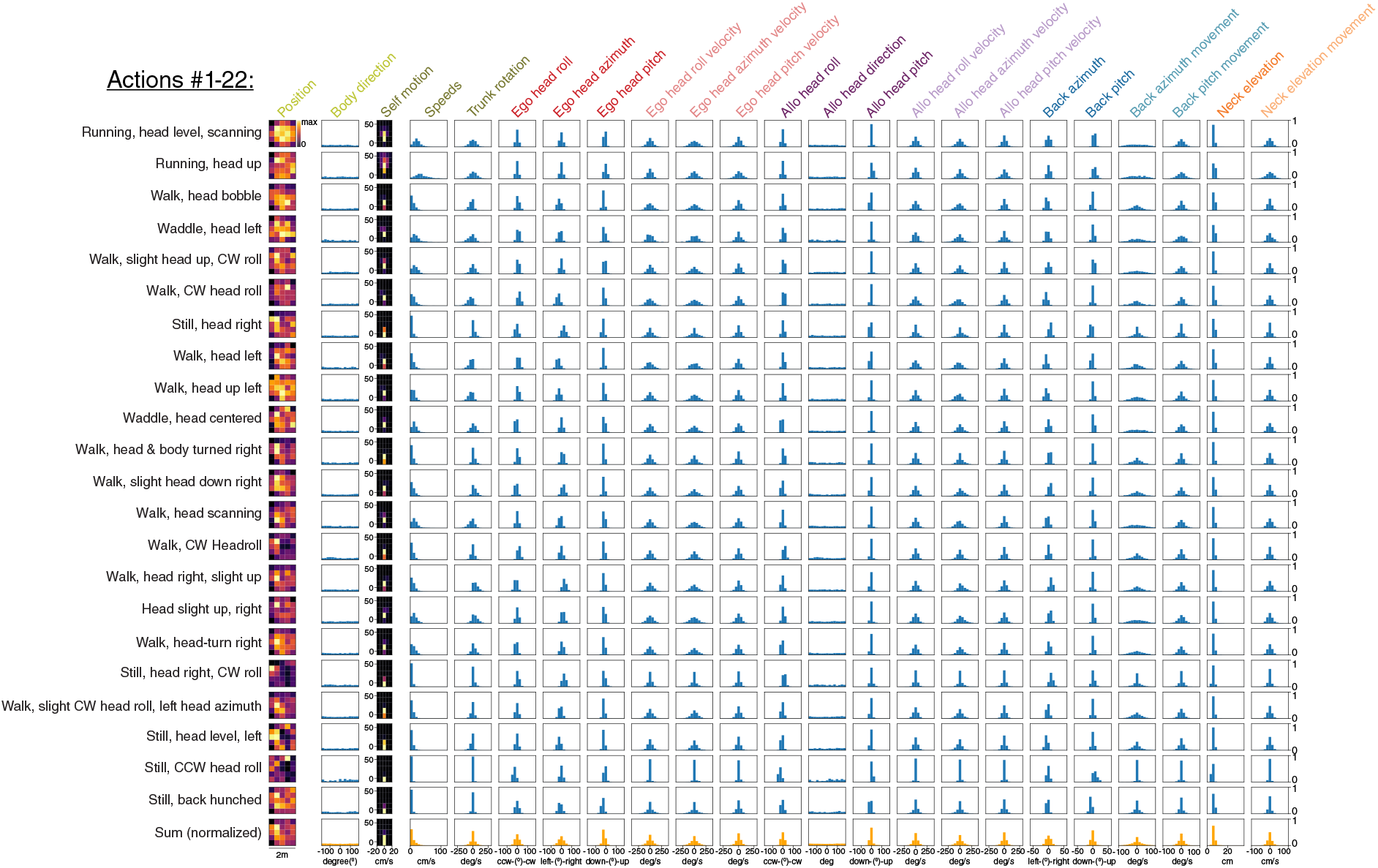
Elementary kinematic composition of each of the 44 defined actions. Distributions of 23 spatial, postural and movement variables (see methods) for each of the 44 actions (22 in (**A**) and 22 in (**B**)), identified by the descriptor on the left. 2D variables (position and self motion) are expressed in 5×5 bins, and 1D variables in 15 bins. Row 23 in (**A**) and (**B**) displays sample distributions of variables for all actions summed together.

**Fig. S4.**
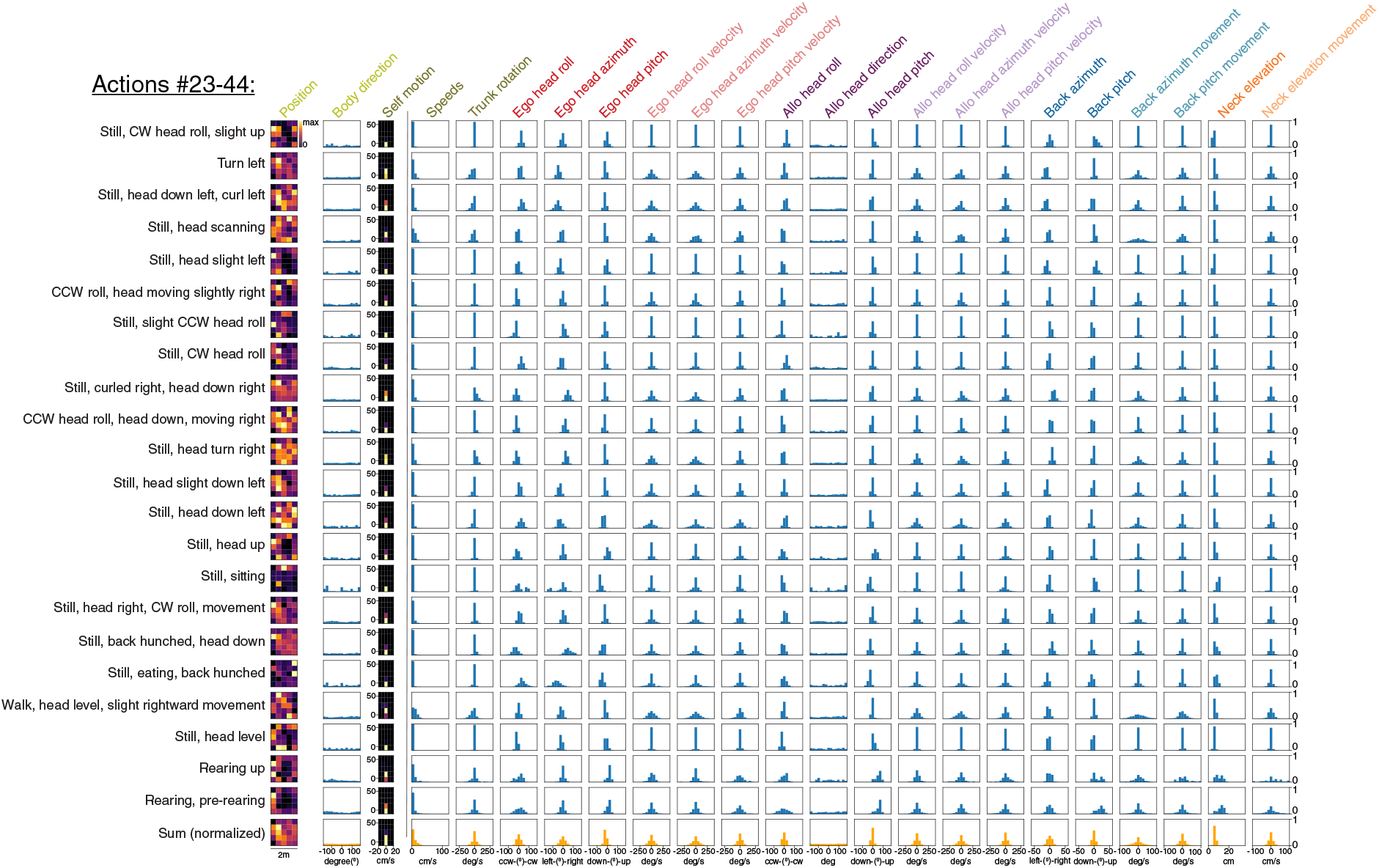
Elementary kinematic composition of each of the 44 defined actions. Distributions of 23 spatial, postural and movement variables (see **methods**) for each of the 44 actions (22 in (**A**) and 22 in (**B**)), identified by the descriptor on the left. 2D variables (position and self motion) are expressed in 5×5 bins, and 1D variables in 15 bins. Row 23 in (**A**) and (**B**) displays sample distributions of variables for all actions summed together.

**Fig. S5.**
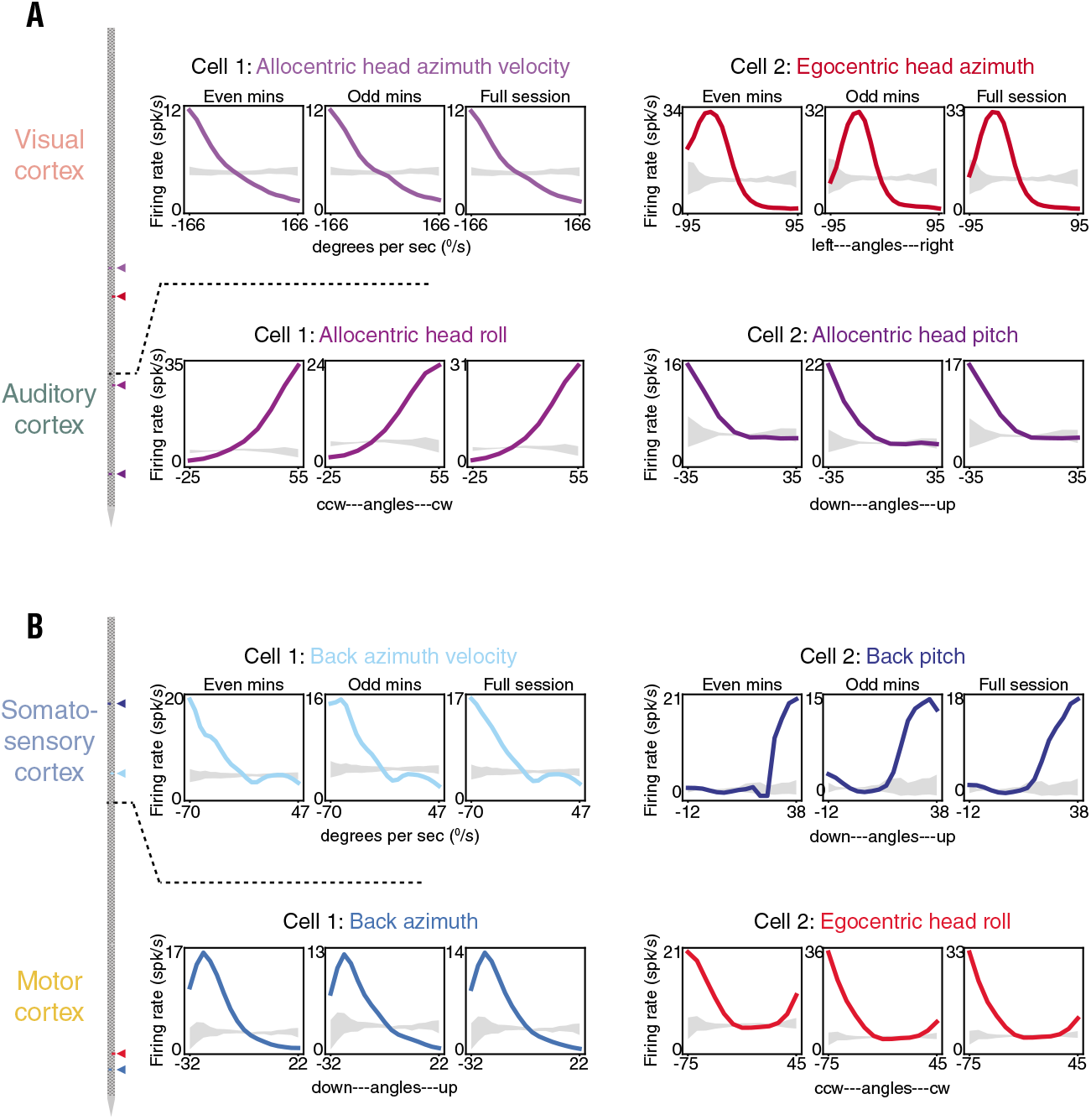
Stable postural and movement tuning curves from each of the four overarching cortical areas. **(A)** (Far left) Recording locations for each neuron are indicated by colored triangles on the Neuropixels schematic. (Top left) Tuning curves from a visual neuron (Cell 1) preferring left-orienting movement of the head in allocentric coordinates (this and all other example cells were recorded in darkness). Data from even and odd minutes of a 20 min recording session are shown adjacently (left and middle) and data from the full session are shown to the right. The 99% CI of the shuffled data are shown in grey. (Top right) Stable tuning curves from a visual cortical neuron preferring left head azimuth in egocentric coordinates (*i.e.*, relative to the trunk). (Lower left) An auditory neuron showing stable tuning to rightward head roll, and (lower right) another auditory neuron preferring downward pitch of the head (in allocentric coordinates). **(B)** A somatosensory cortical neuron tuned to leftward flexion velocity of the back (top left) and an S1 neuron encoding upward pitch of the back (top right). (Lower left) Tuning curves from motor neurons preferring leftward flexion of the back, and (lower right) egocentric head roll to the left.

**Fig. S6.**
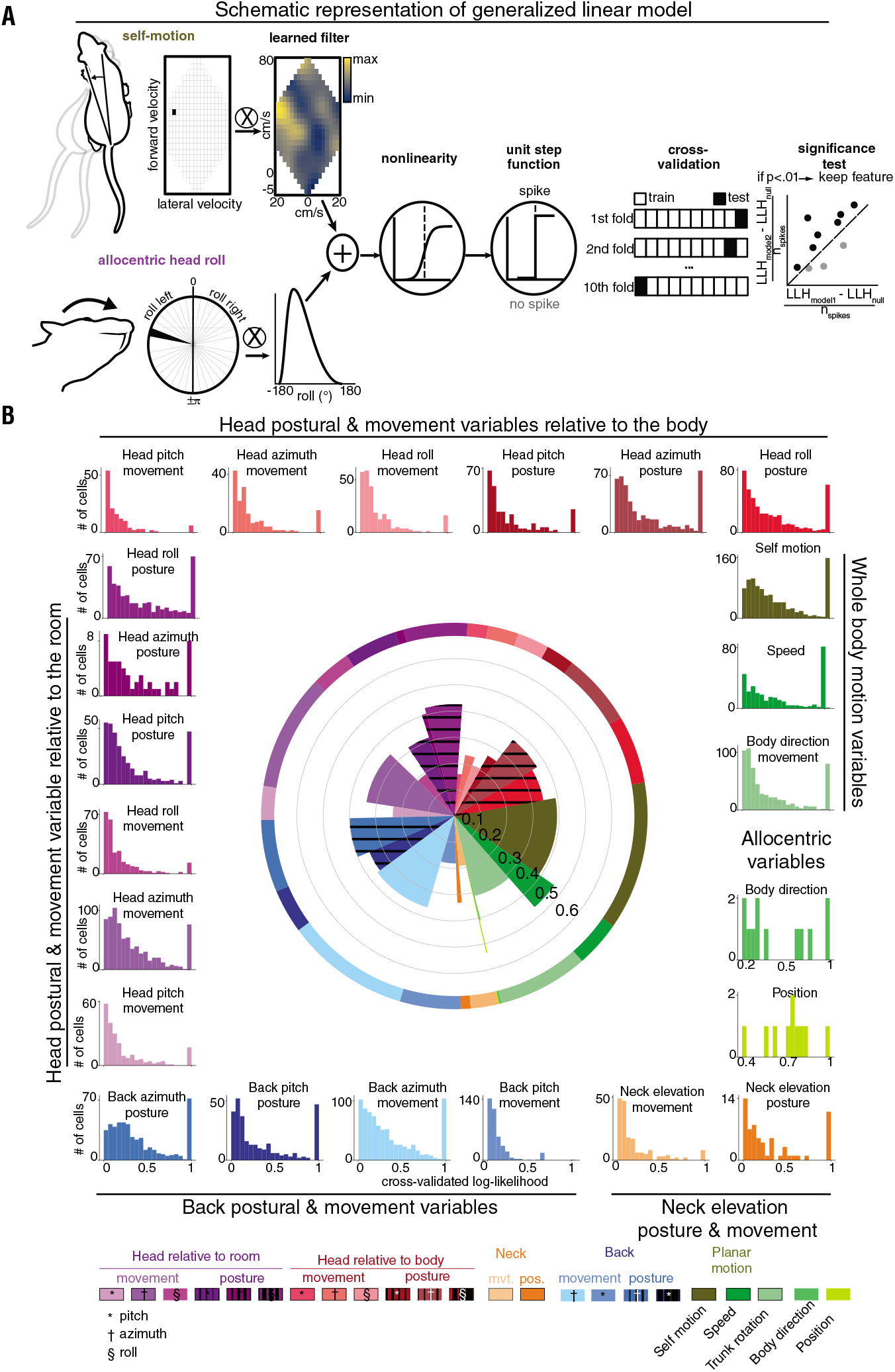
Model selection schematic and contribution to model performance of individual features. **(A)** A toy example of the model selection procedure. A model containing one feature (self-motion, above) is augmented by the addition of a second feature (allocentric head roll, below), as the joint model significantly increases the mean cross-validated rLLR (methods) in explaining the spiking activity of the toy neuron (far right). **(B)** (Middle) Taking all non-null models from all cortical areas together, the width of each wedge indicates the relative share each variable has across all final models and length denotes the mean cross-validated rLLR of that feature across the set of such models. (Outer rim) The distribution of the cross-validated rLLRs for each feature, where the distribution of means correspond to wedge heights and the sum of each histogram normalized by the sum of all histograms corresponds to wedge widths.

**Fig. S7.**
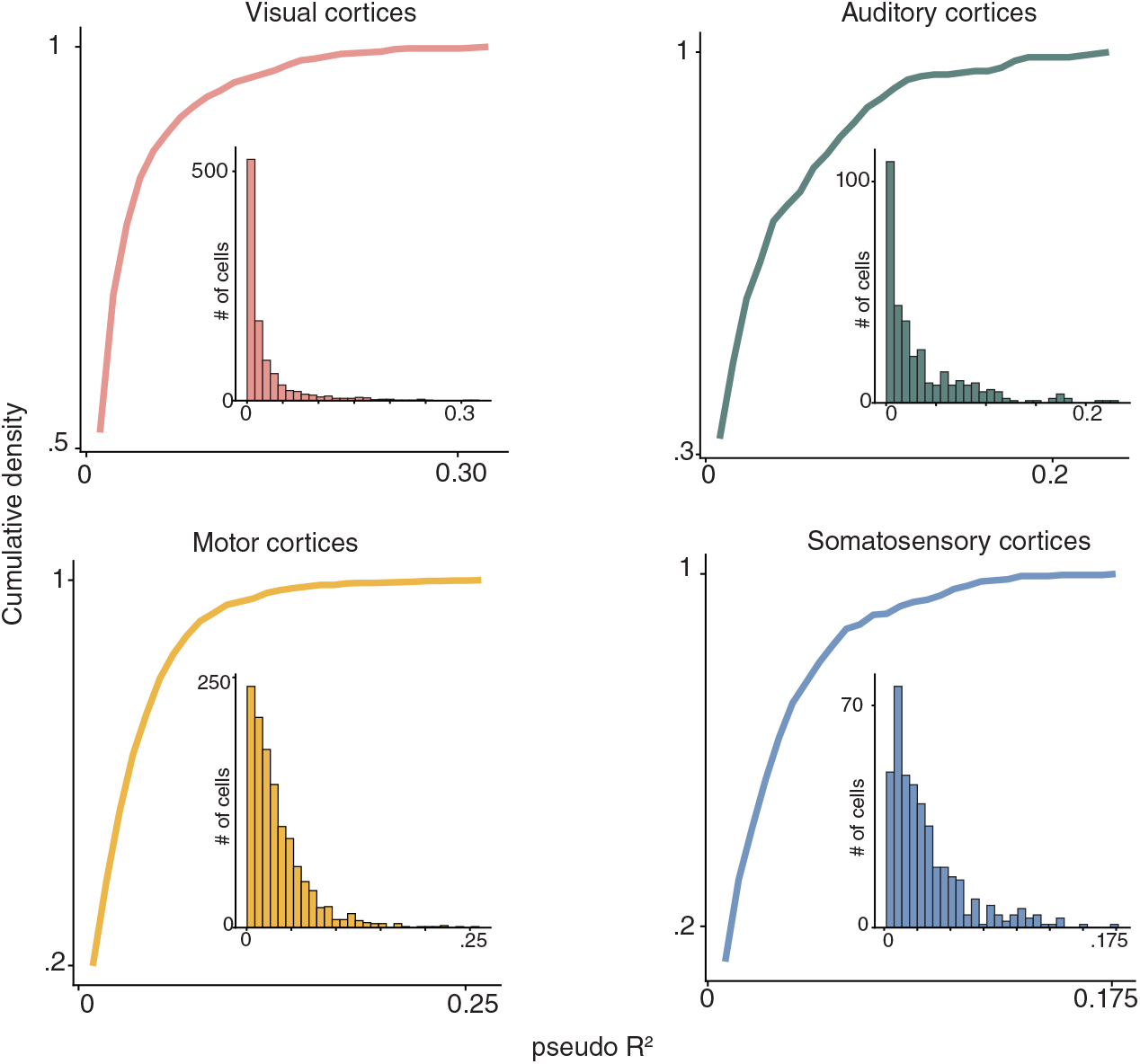
Pseudo-R^2^ of generated statistical models across cortical regions. Mean out-of-sample effect size (McFadden’s pseudo-R^2^) distributions for models generated across the four cortical areas (visual (pink), auditory (cyan), motor (yellow) and somatosensory (blue)).

**Fig. S8.**
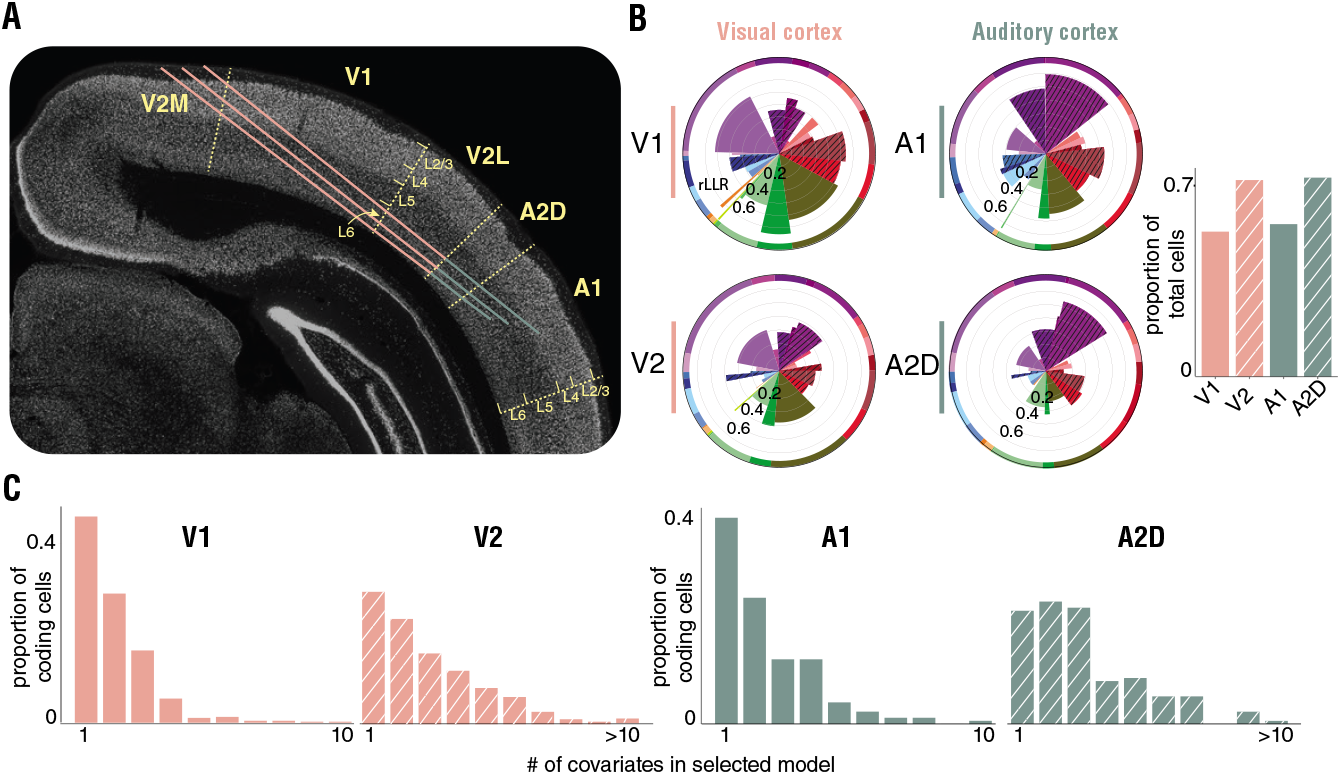
Lamination with comparison of tuning properties across primary and secondary sensory regions. **(A)** Three recording probes plotted against a reference tissue section, displaying the putative cortical layers where single units were recorded in visual and auditory areas. **(B)** (Left) Relative variable share and mean cross-validated rLLR of each feature across models compared across V1 (top) and V2 (V2M, V2L) (below). (Middle) same as (left), but for A1 and A2D. (Right) The proportion of single units encoding at least one behavioral feature in each subregion. **(C)** The proportion of tuned neurons statistically linked to one or any larger number of behavioral covariates (from left to right: V1, V2, A1, A2D).

**Fig. S9.**
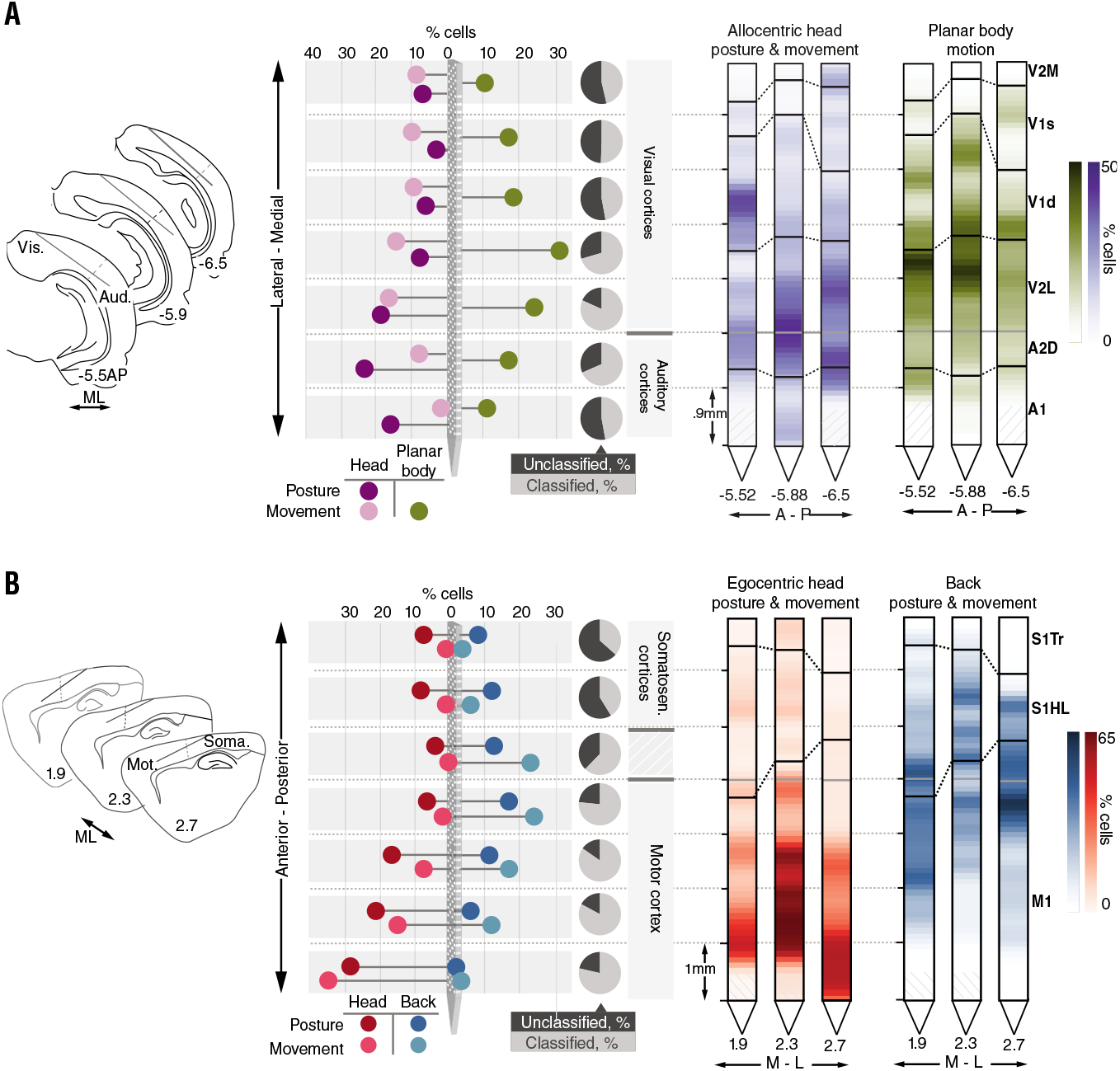
Anatomically organized behavioral tuning gradients. **(A)** (Left) Probe positions relative to corresponding atlas coronal sections in all right hemisphere implanted animals; the dotted line marks the border between auditory and visual cortices. (Middle) Percentage of all recorded cells responsive to allocentric head posture, allocentric head movement and planar body motion along the mediolateral axis. The fraction of cells showing any type of behavioral tuning at a given location is indicated by the black and grey pie charts. (Right) same as (middle), but using the probe view for each animal separately. **(B)** (Left) Same as (A), only for left hemisphere implanted animals, with the dotted line marking the border between primary somatosensory and motor cortices. (Middle) percentage of all recorded cells responsive to egocentric head posture and movement, and back posture and movement along the rostrocaudal axis. (Right) same as (middle), but using the probe view for each animal separately.

**Fig. S10.**
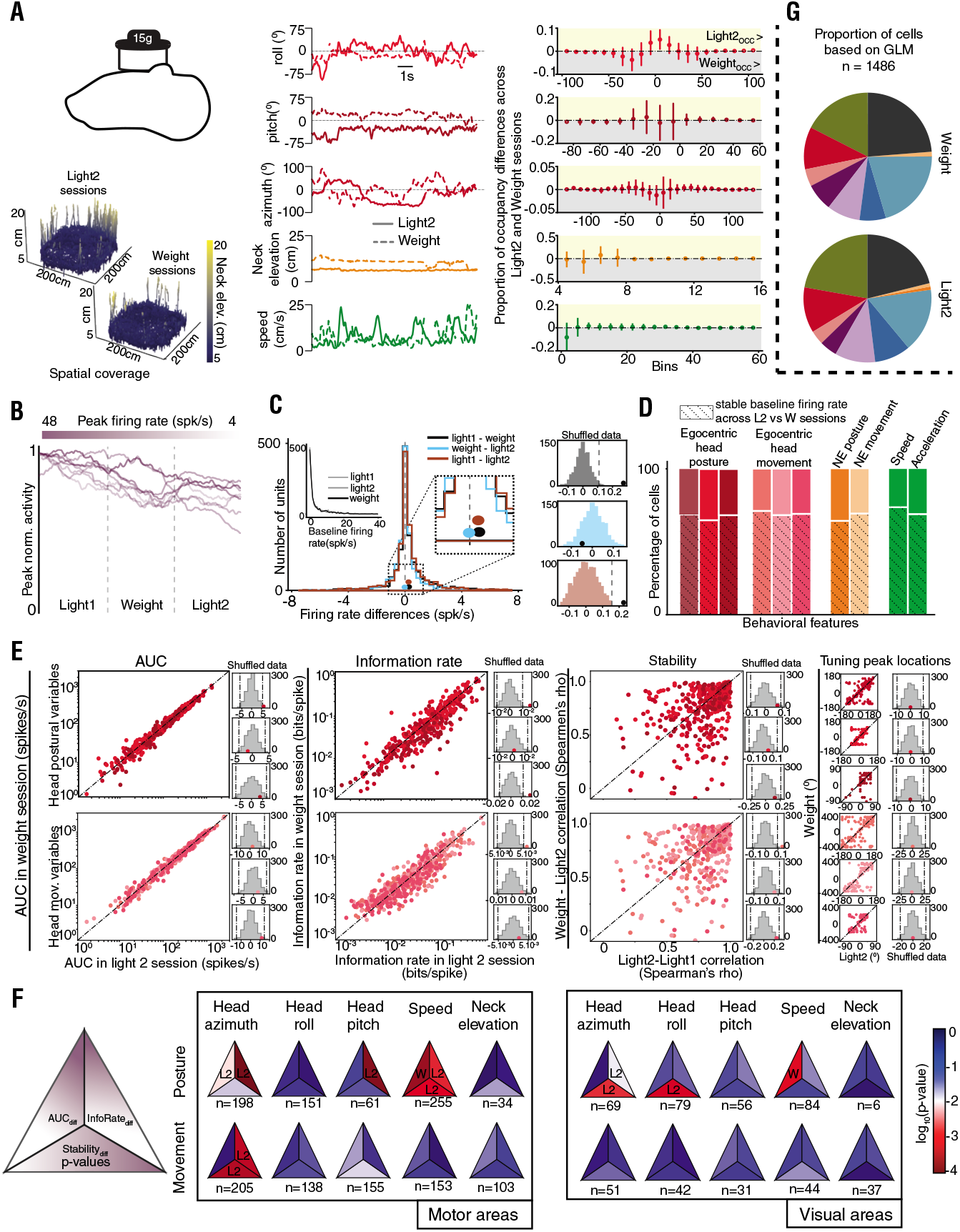
Effects of additional head-mounted weight on behavioral tuning. **(A)** (Upper left) Small, cuboid copper alloy (15 g) relative to the chronic implant on a rat’s head. (Lower left) Differences in the vertical behavior between four left hemisphere (LH-implanted) animals across light/weight sessions. (Middle) Example traces for five features (head azimuth, head roll, head pitch, neck elevation and speed) across light/weight conditions of one LH rat (#26504). (Right) 99% CI of occupancy differences for the same five features across light/weight conditions calculated over all LH and RH rats. **(B)** Firing rate changes across three recording sessions (light1/weight/light2) of seven example single units. **(C)** (Left inset) Baseline firing rate distributions (calculated over the entire session) for all motor single units recorded across light1/weight/light2 sessions. (Middle) Distributions of within-group firing rate differences for motor single units (light1-weight (black), weight-light2 (cyan), light1-light2 (brown)) with distribution means superimposed and magnified (right inset). (Right) Means of each distributions (black dots) relative to their respective shuffled distribution. **(D)** Percentage of significantly tuned cells with a stable baseline firing rate across weight and light2 sessions, relative to all significantly tuned single units in primary motor cortex. **(E)** (Left) Area under the curve, (middle left) information rate, (middle right) stability and (right) tuning peak locations across weight and light2 sessions for all primary motor cortex single units with stable baseline firing rates. The means of the distributions for the three head postural/movement features (red dots) are depicted relative to their respective shuffled distributions (grey). **(F)** (Left) Schematic of what p-value in middle/right is presented in what part of the triangle, “diff” is the comparison between light2 and weight sessions. (Middle) Statistical significance (color) and direction (sign in triangle) of the difference relative to its shuffled distribution for all tuned single units with stable baselines in the primary motor cortex. (Right) Same as middle, but for single units from visual areas. **(G)** GLM results of all primary motor cortex single units across weight and light2 sessions.

**Fig. S11.**
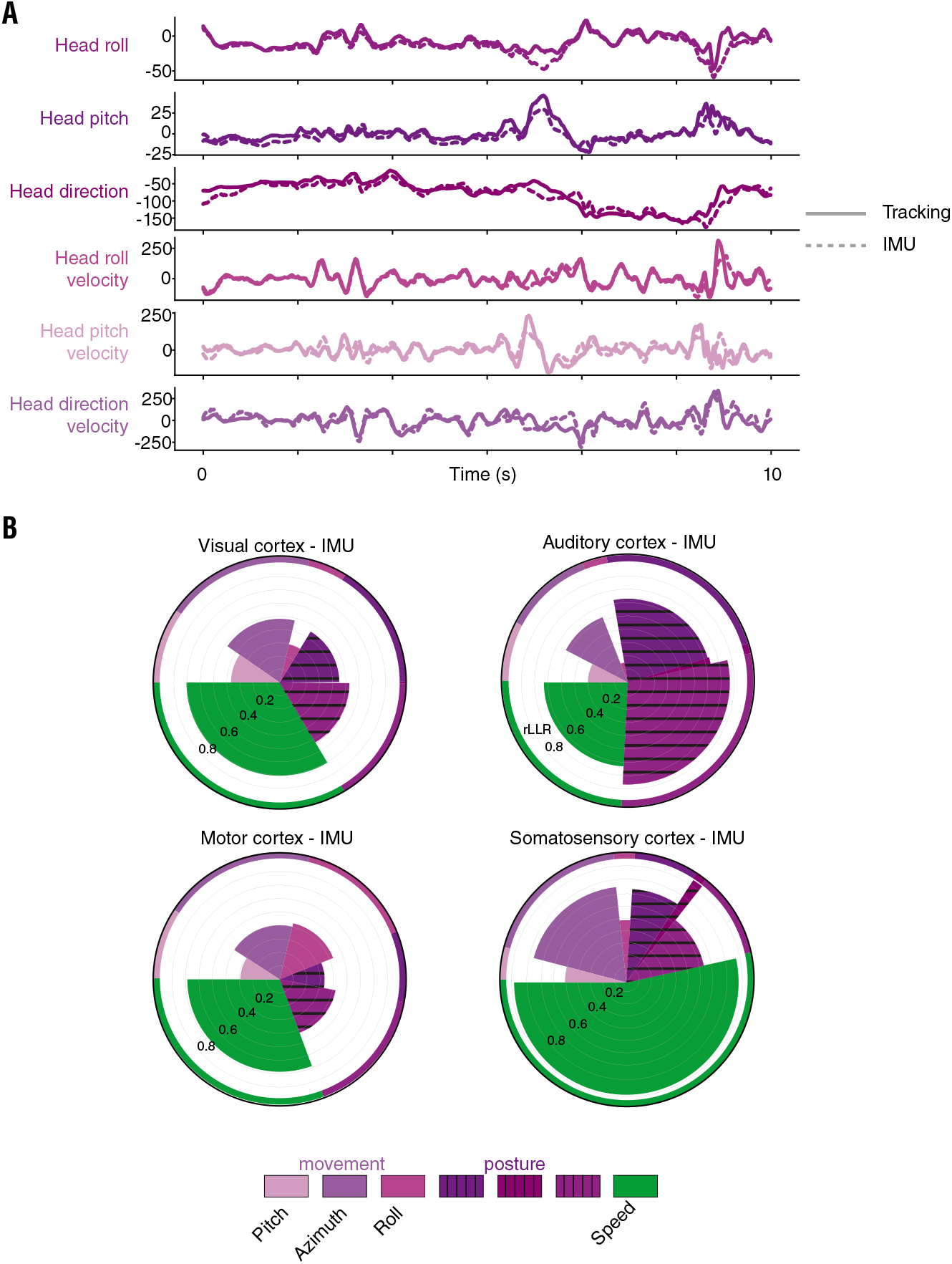
Differences between optical tracking and IMU-generated Euler angles in explaining spiking activity of individual neurons. **(A)** A behavioral recording segment depicting dynamics of six features (allocentric head roll, -head pitch, -head direction, and their velocities), as defined by optical tracking (solid line) or the IMU (dashed line) in one right hemisphere-sampled rat (#26525). **(B)** The relative importance of individual covariates (features in **(A)** and speed) in the data as in **Fig. 2**. (Top) Polar plots represent GLM covariate prevalence from one right hemisphere animal (#26525) using IMU-generated Euler angles for visual cells (left) and auditory cells (right). (Bottom) Polar plots represent GLM covariate prevalence from one left hemisphere animal (#26472) using IMU-generated Euler angles for motor cells (left) and somatosensory cells (right).

**Fig. S12.**
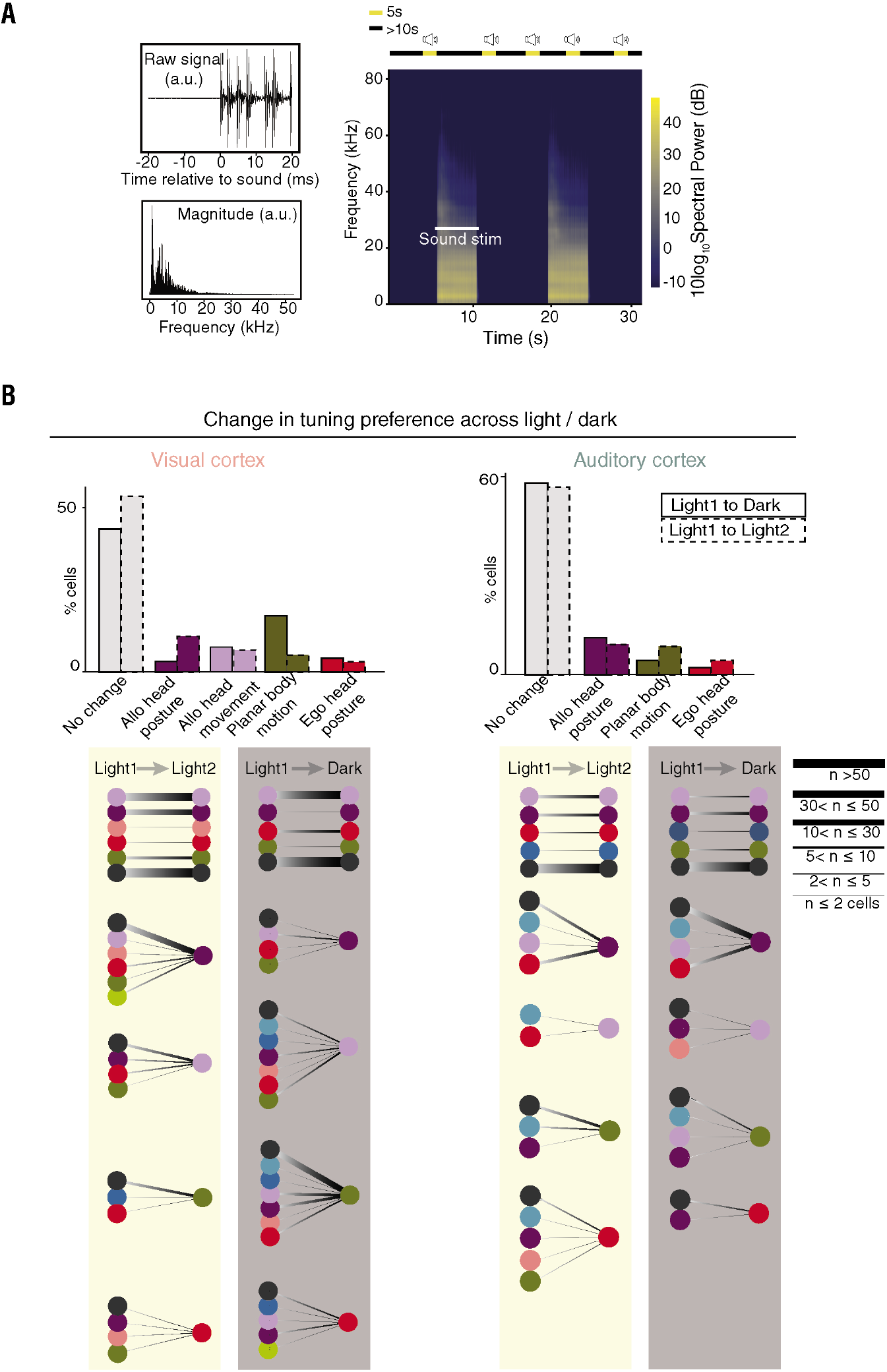
Characteristics of auditory stimuli to assess sound modulation, and comparison of luminance modulation on behavioral tuning in visual and auditory regions. **(A)** (Upper left) Sound amplitude before and during a 20 ms sequence of stimulation. (Lower left) Magnitude of power across the frequency range. (Upper right) Intermittent sound stimulation paradigm during recordings. (Lower right) Sound spectrogram for two example stimulations. **(B)** (Top left) The number of cells showing no change or a change of a certain type between different (light1-dark) and similar (light1-light2) luminance conditions. (Bottom left) The specific types of changes exhibited across different and similar luminance conditions. (Right) Same as (left), only for auditory areas.

**Fig. S13.**
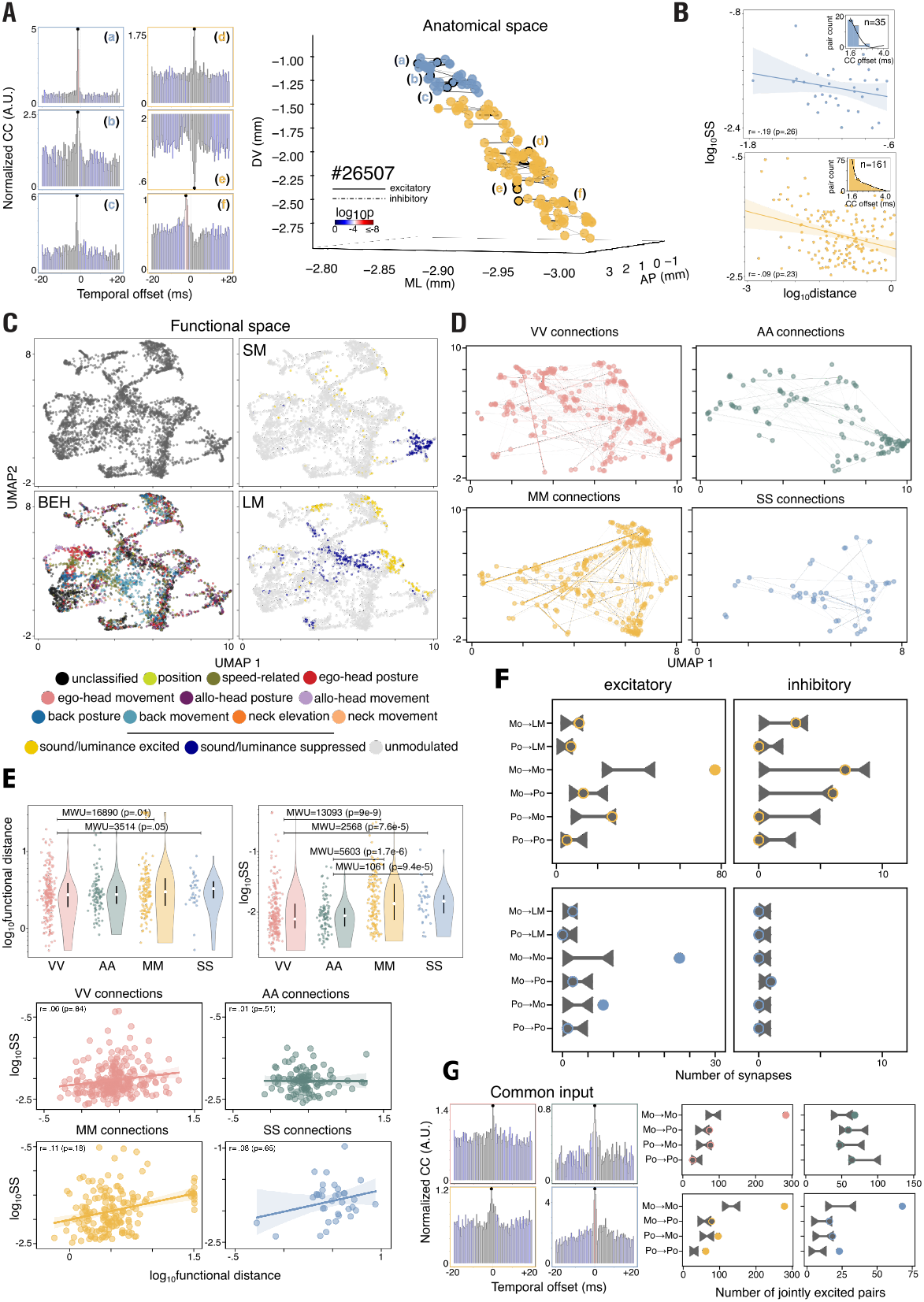
Properties of synaptically coupled neuronal pairs. **(A)** (Left) Three example temporal spiking cross-correlograms from primary somatosensory (a-c) and motor cortices (d-f); (right) all putative synaptic connections including outlined examples (a-f) from one example rat’s (#26507) somatosensory (pink) and motor (green) cortices localized in anatomical space, with the width of each connection weighted by synaptic strength (SS). **(B)** (Top) The log-log relationship between anatomical distance and synaptic strength (SS) for all putative connections in primary somatosensory cortex, (inset) distribution and median (colored circle) of cross-correlogram peaks/troughs; (bottom) same as above, only for primary motor cortex. **(C)** (Top left) All recorded units projected on the low-dimensional covariate embedding; (bottom left) same as (top left), where each single cell is colored by the covariate group that provided the single best fit to its spiking activity. (Top right) Same as (top left) only each single cells is colored by its sound modulation index category. (Bottom right) Same as (top left) only each single cells is colored by its luminance modulation index category. **(D)** (Top left) All visual-visual synaptic connections plotted in functional space (the width of the line denotes synaptic strength). (Bottom left) Same as (top left), only for motor-motor synaptic connections. (Top right) Same as (top left), only for auditory-auditory synaptic connections; (bottom right) same as (top left), only for somatosensory-somatosensory synaptic connections. **(E)** (Top left) Functional distance distributions for all synaptically coupled pairs in the four overarching areas (visual, auditory, motor, somatosensory). (Top right) Same as (top left), only for synaptic strength. (Bottom) Correlations between functional distance and synaptic strength for all four overarching areas (all plots are visualized on log or log-log scales, but all statistics were performed on original data using the Mann–Whitney U (MWU) test). **(F)** Shuffled connection distributions (horizontal line with carets) and experimentally observed excitatory and inhibitory connections (circles) for various functional subtypes in motor (above) and somatosensory (below) cortices. **(G)** (Left) One example cell pair receiving common input for each recorded area (top left - visual, top right - auditory, bottom left - motor, bottom right - somatosensory). (Right) Shuffled common input distributions (horizontal line with carets) and experimentally observed excitatory common input-receiving pairs (circles) for various functional subtypes in visual (top left), auditory (top right), motor (bottom left) and somatosensory (bottom right) cortices.

## Movies

**Movie S1.** Animation of a digitally rendered rat, portrayed by the head, back (3 blue spheres), and neck (smaller green sphere), demonstrating tuning selectivity of a visual cortical neuron, in darkness, to the action “walk, head left”, and a lack of responsiveness for the non-preferred action, “walk, head right”. The session is fast-forwarded between bouts of the specified action and slowed down when the animal is within-behavior; within-behavior bouts are indicated by a green square, both in this and subsequent videos. The translational components of movement have been removed for visualization purposes, though the animals are freely moving.

**Movie S2.** Same as previous movie, but for a neuron in auditory cortex firing selectively during the action “walk, clockwise head roll”; the non-preferred action is “running, head level”.

**Movie S3.** Example of a motor cortical neuron tuned to the action “hunched, head left”, but not responsive to “hunched, head right”.

**Movie S4.** A cell from primary somatosensory cortex firing selectively during bouts of “rearing”, but not “running, head up”.

**Movie S5.** Example of a visual cortical neuron recorded in darkness, tuned to the action “back hunch”, but not “head right”.

**Movie S6.** A neuron in auditory cortex firing selectively during “rearing”, but not during the action “running head up”.

